# *Phytophthora infestans* Ago1-bound miRNA promotes potato late blight disease

**DOI:** 10.1101/2020.01.28.924175

**Authors:** Xinyi Hu, Kristian Persson Hodén, Zhen Liao, Fredrik Dölfors, Anna Åsman, Christina Dixelius

**Author notes:** These authors contributed equally to this work (Xinyi Hu, Kristian Persson Hodén). These authors contributed equally to this work (Anna Åsman, Christina Dixelius). Authors for correspondence: Christina Dixelius, Anna Åsman.

## Abstract

- *Phytophthora* spp. incite serious plant damages by exploiting a large number of effector proteins and small RNAs (sRNAs). Several reports are describing modulation of host RNA biogenesis and defence gene expression. Here, we analysed *P. infestans* Argonaute (Ago) 1 associated small RNAs during potato leaf infection.
- sRNAs were co-immunoprecipitated, deep sequenced and analysed against the *P. infestans* and potato genomes, followed by transgenic and biochemical analyses on a predicted host target.
- Extensive targeting of potato and pathogen-derived sRNAs to a large number of mRNAs was observed, including 206 sequences coding for resistance (R) proteins in the host genome. The single miRNA encoded by *P. infestans* (miR8788) was found to target a potato lipase-like membrane protein-encoding gene (*StLL1*) localized to the tonoplast. Analyses of stable transgenic potato lines harbouring overexpressed *StLL1* or artificial miRNA gene constructs demonstrated the importance of StLL1 during infection by *P. infestans*. Similarly, a miR8788 knock-down strain showed reduced growth on potato compared to the wild-type strain 88069.
- The data suggest that sRNA encoded by *P. infestans* can affect potato mRNA and thereby promote disease. Knowledge of the impact of pathogen small RNAs in plant defence mechanisms is of major significance to succeed in improved disease control management.

## Introduction

Gene regulatory mechanisms in eukaryotes are complex involving large number of proteins, regulating nucleosome and chromatin modifications and involvement of a range of transcription factors. Small non-coding RNAs (sRNAs) have added new insights into these processes not least on post-transcriptional and post-translational regulation. sRNAs are known to trigger sequence-specific cleavage or translational repression of target transcripts. sRNAs are well studied in several model species including Arabidopsis, since they are involved in numerous processes including development and stress responses (Axtell, 2013). In plants, 21-22 nucleotide (nt) sRNAs are processed from RNA polymerase II-transcribed primary RNAs (microRNAs or miRNAs) or generated from dsRNA. Among the latter group, secondary small interfering siRNAs (siRNAs) can be formed such as phasiRNA and tasiRNAs. Whereas the 24-nt siRNAs (p4-siRNAs) predominately expressed in developing endosperm is treated as a third main group of sRNAs. In the centre for all these processes are Dicer-like (DCL), Argonaute (AGO) and RNA dependent RNA polymerase (RDR) protein families. There are however distinct differences in sequence formation and preferences, biogenesis and downstream processing of the different sRNA categories. Additional proteins take part in transcriptional and post-transcriptional gene silencing events that together with the different sRNA classes form a complex picture of the gene regulatory processes, which are excellently reviewed elsewhere (Rogers & Chen, 2013; Matzke & Mosher, 2014; Borges & Martienssen, 2015; Budak & Akpinar, 2015; Fang & Qi, 2016).

Besides endogenous gene regulation of resistance (*R*) genes particularly by miRNAs (Shivaprasad *et al.*, 2012; Yang *et al.*, 2013; Budak *et al.*, 2015), much emphasis has been turned to the mobility of sRNAs within or between different organisms. Initially, RNA interference in plants was devoted to studies of virus responses. It was shown that virus infection is arrested by producing DCL-dependent and virus-derived siRNAs which guide plant AGO proteins to viral RNAs (Guo *et al.*, 2019). To subvert plant defence responses, eukaryotic pathogens can deliver sRNAs during infection such as shown by the fungus *Botrytis cinerea* (Weiberg *et al.*, 2013). In this case the pathogen hijacks the sRNA machinery in Arabidopsis with its own sRNAs. However, there are also cases where for example plant miRNAs (miR159 and miR166) can be induced upon fungal infection and cleave fungal effector mRNAs resulting in resistance to the fungus *Verticillium dahliae* (Zhang *et al.,* 2016). Although the number of reported cases of plant-pathogen exchange of sRNAs is increasing, there is a lack of deeper understanding of RNA-based events between crop species and their major disease causing agents.

The late blight disease of potato (*Solanum tuberosum*) and tomato (*S. lycopersicum*) incited by the oomycete *Phytophthora infestans* causes global annual losses estimated to € 12 billions (Arora *et al.*, 2014). *P. infestans* has a large genome enriched in genes coding for effector proteins that promote plant infection by secretion into the apoplastic and cytoplasmic spaces of the host tissue (Haas *et al.*, 2019; Whisson *et al.*, 2016). These virulence genes are predominantly located among repeat sequences, explaining the rapid adaptation potential of this pathogen. Constant changes of *P. infestans* populations and emergence of new aggressive strains make disease control including resistance breeding extraordinary challenging. Most of the gene silencing components found in eukaryotic organisms are present in *P. infestans* (*Pi*), such as proteins encoded by two *DCL* genes (*PiDcl1*-*2*), five *AGOs* (*PiAgo1-5*) and one gene coding for RDR, *PiRdR1* (Vetukuri *et al.*, 2011; Fahlgren *et al.*, 2013). Silencing-related proteins such as cytosine methyltransferases including HUA Enhancer 1 (HEN1), RNA polymerase IV, Drosha and ERI1, present in other eukaryotic organisms, are however not identified in its genome (Vetukuri *et al.*, 2011). Adenine N6-methylation (6mA) has in case of *P. infestans* and *P. sojae* replaced the more common 5-methylcytosine DNA-methylation implicated in gene regulation (Chen *et al.*, 2018). New functional pathways and or new candidates being involved in canonical pathways need to be clarified. The potato genome is large (Xu *et al.*, 2011; Hardigan *et al.*, 2016) which hampers detailed analysis of sRNAs and their predicted targets. So far are 21-24 nt sRNAs reported with the 24 nt as the major size class (Lakhotia *et al.*, 2014). The number of components taking part in the sRNA biogenesis, their function and responses induced upon pathogen attack remain to be demonstrated.

Here, we used an *Ago1-GFP P. infestans* strain (Åsman *et al.,* 2016) to analyse sRNA-associated events during potato infection. We show extensive targeting of potato and pathogen-derived sRNAs to a large number of mRNAs, including 206 sequences coding for resistance (R) proteins in the host genome. Intriguingly, the single miRNA in *P. infestans* (miR8788) was found to target a lipase-like membrane protein-encoding gene *StLL1* whose suppression promotes pathogen growth. miR8788 seems to be an ancient molecule, tracing back to strains from 1846, which highlights its conserved role in abating the defence machinery to this devastating pathogen.

## Materials and Methods

### Materials for Ago-RNA co-immunoprecipitation (co-IP)

Two transformants of *P. infestans* wildtype (WT) strain 88069 were used (Åsman *et al.*, 2016): one harboring a green fluorescent protein-encoding gene (*pHAM34:eGFP*), and the other *P. infestans Ago1-GFP* (*pHAM34:PiAgo1-GFP*). Plant growth conditions, pathogen storage, cultivation and inoculation procedures were as earlier described (Vetukuri *et al.*, 2011; Jahan *et al.*, 2015). Three replicates of leaves from cv. Bintje inoculated with either of the two *P. infestans* strains were collected 6 days post inoculation. For each replicate, 2 g of leaf material was pooled (Fig. S1). Mycelia of the two *P. infestans* strains were grown in pea broth for 7 days, before collecting 3 mycelia replicates of 200 mg from each strain.

### RNA co-IP, sequencing and data analysis

IP of the collected leaf materials and mycelia was performed (Åsman *et al.*, 2016). Twelve libraries were made at the Ion Proton platform, SciLifeLab (Uppsala, Sweden), where the Ion PI Sequencing 200 kit v3 (Life Technologies) was used. Reads of length 18-38 nt were adaptor-trimmed and mapped to the *P. infestans* genome (Haas *et al.*, 2009) and the *Solanum tuberosum* genome v4.04 (Hardigan *et al.*, 2016) using Bowtie2 v. 2.3.2 (Langmead & Salzberg, 2012). Reads from ribosomal RNA (rRNA) and transfer RNA (tRNA) were discarded. Remaining sRNAs (duplicates excluded) were counted using Shortstack v3.8.2 (Johnson *et al.*, 2016). A cut-off of 10 RPM (reads per million) was set for inclusion of sRNA in the analysis. For downstream analysis, an enrichment of at least 20% RPM compared to the control (GFP) was considered significant (miR8788 was detected only in the PiAgo1 co-IP data). No mismatches were allowed and all mapping locations were reported. *P. infestans* and potato annotations were downloaded from http://protists.ensembl.org/ and http://solanaceae.plantbiology.msu.edu/. Reads mapping to miRNA were predicted using Shortstack, with following settings --strand_cutoff 0.5 --foldsize 1000 --dicermin 18 --dicermax 38. For pipeline details, see Fig. S2. PhasiRNAs were predicted using PhaseTank v1.0 (Guo *et al.*, 2015).

### sRNA target prediction

To predict sRNA targets in potato, the predicted sRNA from Shortstack was run in the target analysis server psRNATarget (Dai *et al.*, 2018), against the cDNA library of “*S. tuberosum* transcript, JGI genomic project, Phytozome 12, 448_v4.03” with default settings, except the expectation value was 3. The miR8788 target prediction was run with expectation value 5 (default). sRNA targets in *P. infestans* were predicted using the sRNA from Shortstack processed in TargetFinder (https://github.com/carringtonlab/TargetFinder) against the *P. infestans* genome (Haas *et al.*, 2009), with default settings.

### Multiple sequence alignment and phylogeny

100 homologs with the highest similarity score compared to StLL1 or AtLL1, respectively, were collected from NCBI (2018-04-15) using BLASTP taxonomy function with default settings. Additional solanaceous sequences were mined from https://solgenomics.net/, http://www.onekp.com, http://www.ebi.ac.uk (PMID:26442032) and http://www.doi.org/10.5061/dryad.kc835.5. cDNA from *Calibrachoa hybrida* was amplified, sequenced (MK550718) and added to the generated sequence dataset. Single-gene maximum likelihood (ML) trees were performed on RAxML v.8.2.11 (Stamatakis, 2007) using the PROTGAMMAJTT and autoMRE settings, including 200 bootstrap replicates. The tree was depicted using the R package ggtree and positive selection was tested using PAML v. 4.9h (Yang, 2007).

### *StLL1* cloning, transient expression in *Nicotiana benthamiana*, Western blot and confocal microscopy

Total RNA was isolated from potato using the RNeasy plant mini kit (Qiagen) and cDNA was synthesized with the Maxima cDNA synthesis kit (Thermo Fisher Scientific) throughout the work. The *StLL1* CDS was cloned from potato cv. Sarpo Mira and fused in the 3’ end to *GFP*, in the binary plasmid pGWB505 (Nakagawa *et al.*, 2007) and transformed into *Agrobacterium* strain GV3101 containing pSoup (Hellens *et al.*, 2000). The *P*. *infestans* miR8788 stem-loop sequence was amplified from *P*. *infestans* strain 88069 genomic DNA. The complete stem-loop structure was cloned (Gateway, Invitrogen Life Technologies) into pGWB505, forming a *p35S*:*miR8788*-*GFP* construct. All constructs generated in this study were confirmed by Sanger sequencing (Macrogene Inc.), and all cloning primers are listed in Table S1. Three-week-old *N. benthamiana* plants were used for Agro-infiltration. Samples of leaves co-infiltrated with *StLL1-GFP* and the stem-loop of either miR8788 or ame-miR6055 (miR6055) were collected after three days and used for Western blot analysis (Åsman *et al.*, 2016), using mouse monoclonal anti-GFP antibody (JL-8, BioNordika, In Vitro Sweden AB). Six-week-old *N. benthamiana* plants were Agro-infiltrated with pGWB505 containing *p35S*:*StLL1*-*GFP* and the p19 silencing suppressor (Scholthof, 2006) and used for sub-cellular localization analysis. Control leaves were infiltrated with p19 alone. Staining of the plasma membrane was done by soaking leaf pieces in 50 µM FM4-64FX (Thermo Fisher Scientific) for 30 min. Imaging was performed 3-5 days post infiltration using an LSM800 confocal microscope (Zeiss). Excitation/detection wavelengths for GFP and FM4-64FX were 488/410-546 nm and 506/650-700 nm, respectively. FM4-64FX (red) colour was changed to magenta for colour-blind visibility in ImageJ. Brightness and contrast were increased with the same amount for all images (6-175 on histogram).

### Transient transcription dual-luciferase (Dual-LUC) assays

A transient transcription Dual-LUC assay was used (Banerjee *et al.*, 2007; Liu *et al.*, 2008). The U6-26 promoter was amplified from the pHEE401E backbone (Wang *et al.*, 2015) and inserted into the *Kpn*I/*Not*I site of pGreenII8000. After *Not*I and *Xba*I digestion, the *StLL1* target sequence (TS) was ligated to Firefly luciferase CDS. Primer sequences are provided in Table S1. The *StLL1* mutated target sequence (NS) acts as control (Fig. 2). The luciferase activity was analyzed using white 96-well plates (Nunclon), and LUC reaction reagents according to the manufacturer’s instruction (Promega, Dual-Luciferase® Reporter Assay System. The measurements were done on a plate luminometer equipped with dual injectors (TECAN infinite M1000 PRO).

### *P. infestans* miR8788 cloning and 5’RLM-RACE

5’RLM-RACE was performed as follows (McCue et al. 2013): GeneRacer™ RNA Oligo was ligated to the total RNA using T4 RNA ligase (Nordic Biolabs). cDNA was synthesized and PCR-amplified. First, a touch down PCR with GeneRacer™ 5′ primer and a gene-specific reverse primer was run. The GeneRacer™ 5′ nested primer was used for a second PCR where the first PCR products served as template. After gel purification, the amplified products were cloned into pJET1.2/blunt (Thermo Fisher Scientific) and sequenced (Macrogen). The oligo sequences are found in Table S1.

### *P. infestans* miR8788 knock-down

A miRNA target mimicry approach (Franco-Zorrilla *et al.*, 2007) was applied to silence the miR8788 sequence located in the *PITG_10391* gene. GFP-MIM8788a/MIM8788b fusions were generated from the pTOR-NGFP vector (Åsman *et al.*, 2016) using Phusion high-fidelity DNA polymerase (Thermo Fisher Scientific) and a reverse GFP cloning primer that contained a miRNA mimicry sequence (MIM8788a or MIM8788b). The PCR product was digested with *Eco*RI and *Not*I, and the fragment was ligated into the corresponding restriction sites in the pTOR-NGFP vector, thereby replacing NGFP. The final insert was 738 bp (717bp GFP, 21bp MIM8788a/MIM8788b) and driven by the *HAM34* promoter. Transformation of the *P. infestans* 88069 WT strain was performed as earlier outlined (Vetukuri *et al.*, 2012), using 50 μg plasmid DNA for each transformation experiment. RNA was extracted from *P. infestans* (88069, *pHAM34:PiAgo1-GFP* and miR8788 knock-down (KD)) mycelia using Trizol (Thermo Fisher Scientific). One microgram total RNA was reverse transcribed into cDNA using Maxima Reverse Transcriptase (Thermo Fisher Scientific) as earlier described (Chen et al. 2005). The expression of miR8788-5p and −3p was quantified using iTaq Universal SYBR Green Supermix (Bio-Rad) on the CFX Connect Real-Time PCR Detection System (Bio-Rad), using *Pi-Actin A* (PITG_15117) as reference gene.

### Plant transformation

*StLL1* RNAi primers were designed by WMD3 (Web MicroRNA Designer http://wmd3.weigelworld.org/cgi-bin/webapp.cgi). The RNAi vector pRS300 was from Addgene and the *StLL1* amiRNA (*StAmiRNA*) precursor was ligated into pGWB505. The pGWB505 empty vector, pGWB505-StLL1 and pGWB505-amiRNA-StLL1 were transformed into potato as earlier outlined (Jahan *et al.*, 2015). Both Desirée and Sarpo Mira cultivars were used to ensure production of transformed plants. Ten potential transgenic shoots per construct were grown on MS media containing 20 µg hygromycin B/ml. Transgenic plants were inoculated with *P. infestans* (strain 88069 on Desirée and for Sarpo Mira strain 11388) as earlier described (Jahan *et al.*, 2015). The strain 11388 is very aggressive and has overcome the resistance in cv. Sarpo Mira and was used in case of Sarpo Mira infections. The 88069 strain induces disease on cv. Bintje and Desirée. RNA was extracted from *P. infestans* inoculated cv. Sarpo Mira and Desirée leaves, and cDNA was synthesized. Maxima SYBR Green/Fluorescein Master Mix (Thermo Fisher Scientific) was used for the qRT-PCR reactions (BioRad iQ5) and miRNA was quantified (Varkonyi-Gasic *et al.*, 2007), using *Pi-Actin A* (PITG_15117) as reference gene.

### *P. infestans* DNA quantification

To quantify DNA of *P. infestans* in infected leaves, qPCR analysis was carried out essentially as earlier described (Llorente *et al.*, 2010). Genomic DNA from infected potato leaves was isolated using CTAB lysis buffer at 65°C for an hour followed by phenol/chloroform extraction. 20 ng DNA was used in qPCR analysis and at least three biological replicates were analysed. Primers are listed in Table S2.

### Northern hybridization

Total RNA from potato and *P. infestans* (strains 88069, *pHAM34:PiAgo1-GFP* and *pHAM34:GFP*) was isolated using PureLink™ Plant RNA Reagent for plant samples (Thermo Fisher Scientific) and Trizol for mycelia samples (Thermo Fisher Scientific). Nineteen µg RNA enriched for the low-molecular weight fraction (Kreuze *et al.*, 2005) was resolved on denaturing 12.5% polyacrylamide gels. γ-^32^P labelled RNA probes (Sigma) (Table S1) were used in Northern hybridization (Åsman *et al.*, 2016). The miR8788 probes and *P. infestans* 5S rRNA probe were hybridized at 42°C whereas the potato U6 snRNA probe was hybridised at 54°C.

### Statistical analysis

The Levene’s test was used to test for homogeneity of variance across groups (Brown & Forsythe, 1974). When no inequality in variance between samples was found, one of following three approaches implemented with R was run depending on the experimental setup. For pairwise comparisons, Student’s unpaired *t*-test was employed. One-way ANOVA was used when analysing the influence of one variable on the difference in mean between three or more samples. Experiments examining the influence of two different variables on three or more samples were analysed with two-way ANOVA. Correction for multiple comparisons in any ANOVA test was performed with Tukey’s HSD test.

### R packages used in this study

The following R packages were utilized: ggtree (v.1.16.6, https://bioconductor.org/packages/release/bioc/html/ggtree.html), car (v.3.0.3, https://cran.r-project.org/web/packages/car/index.html), onewaytests (v.2.4, https://cran.r-project.org/web/packages/onewaytests/index.html), stats (v.3.6.1, https://www.R-project.org/), userfriendlyscience (v. 0.7.2, https://cran.r-project.org/web/packages/userfriendlyscience/index.html).

### Data Availability

The sRNA sequencing data is available through the National Center for Biotechnology Information (NCBI) Gene Expression Omnibus, series accession number GSE119230. Novel miRNA are available at miRBase upon acceptance. Additional sequence and phylogeny data can be found in the Treebase repository, http://purl.org/phylo/treebase/phylows/study/TB2:S24467.

## Results

### PiAgo1 shifts 5’ nt preference from C to U during infection

We infected potato leaves with the same *P. infestans pHAM34*:*PiAgo1-GFP* strain as in the earlier study, and included as controls leaves infected with a strain containing *pHAM34:eGFP* (Avrova *et al.*, 2008) (Fig. S1) and mycelia from the two strains grown on plates. Co-immunoprecipitated sRNAs were analysed on an Ion Proton sequencing platform, generating in total 86,600,701 raw reads. Computational processing (Fig. S2) resulted in six sRNA datasets comprising 26,123,643 sRNA reads from *P. infestans* and 3,090,568 potato sRNA reads. Analysis of the *P. infestans* sRNA distribution indicated size enrichment of 21 nt sRNAs both in the mycelia sample and in the sample from infected potato leaves. The proportion of 21 nt sRNAs was higher in pure mycelia (42,275 reads, 54%) than in the leaf-infected sample (24,088 reads, 39%) (Fig. 1a, Fig. S3) and there was a slight enrichment of 25 and 26 nt *P. infestans* sRNAs during potato infection. This size pattern is similar to the previously reported information on the highly pathogenic *P. infestans* strain 3928A (Vetukuri *et al.*, 2012). Unexpectedly, potato sRNAs could be detected in our PiAgo1 pull-down material, indicating a potential incorporation of host sRNAs into the oomycete RNAi machinery. The potato sRNA size profile showed enrichment between 18 to 24 nt with a peak at 21 nt and minor presence of sRNAs between 25 to 37 nt (Fig. 1b). Plant miRNAs are known to have a predominant length of 21 nt, particularly from DCL1 processing, while somewhat longer miRNAs are produced by other DCL proteins (Rogers & Chen 2013). Most plant miRNAs have unique 5’-terminal U nucleotides, a feature selected by Ago1 for the gene silencing process (Mi *et al.*, 2008). A clear U preference was found in PiAgo1-associated sRNAs mapping to potato mRNA, particularly among the 21 nt sRNAs (Fig. 1c). The 5’U preference is frequently reported in other organisms, not least in fungi (Thieme *et al.*, 2012; Dahlman & Kück, 2015). The 5’U bias among 21 nt PiAgo1-sRNAs is noteworthy, considering the 5’C preference previously observed among sRNAs of this size class in PiAgo1 at the mycelium stage (Åsman *et al.*, 2016). This shift in PiAgo1 5’ nt preference during infection could be either (i) an indirect consequence of the 5’U bias of plant 21 nt sRNAs (Mi *et al.*, 2008), or (ii) due to selective preference of PiAgo1 for host miRNAs, whereof the vast majority are 21 nt and have 5’U (Mi *et al.*, 2008; Thieme *et al.*, 2012). An intriguing possibility could be that *Phytophthora* Ago1 has co-evolved with host miRNA (size and 5’ nt) as a means to efficiently down-regulate selected host genes to favour infection.

**Figure 1.**
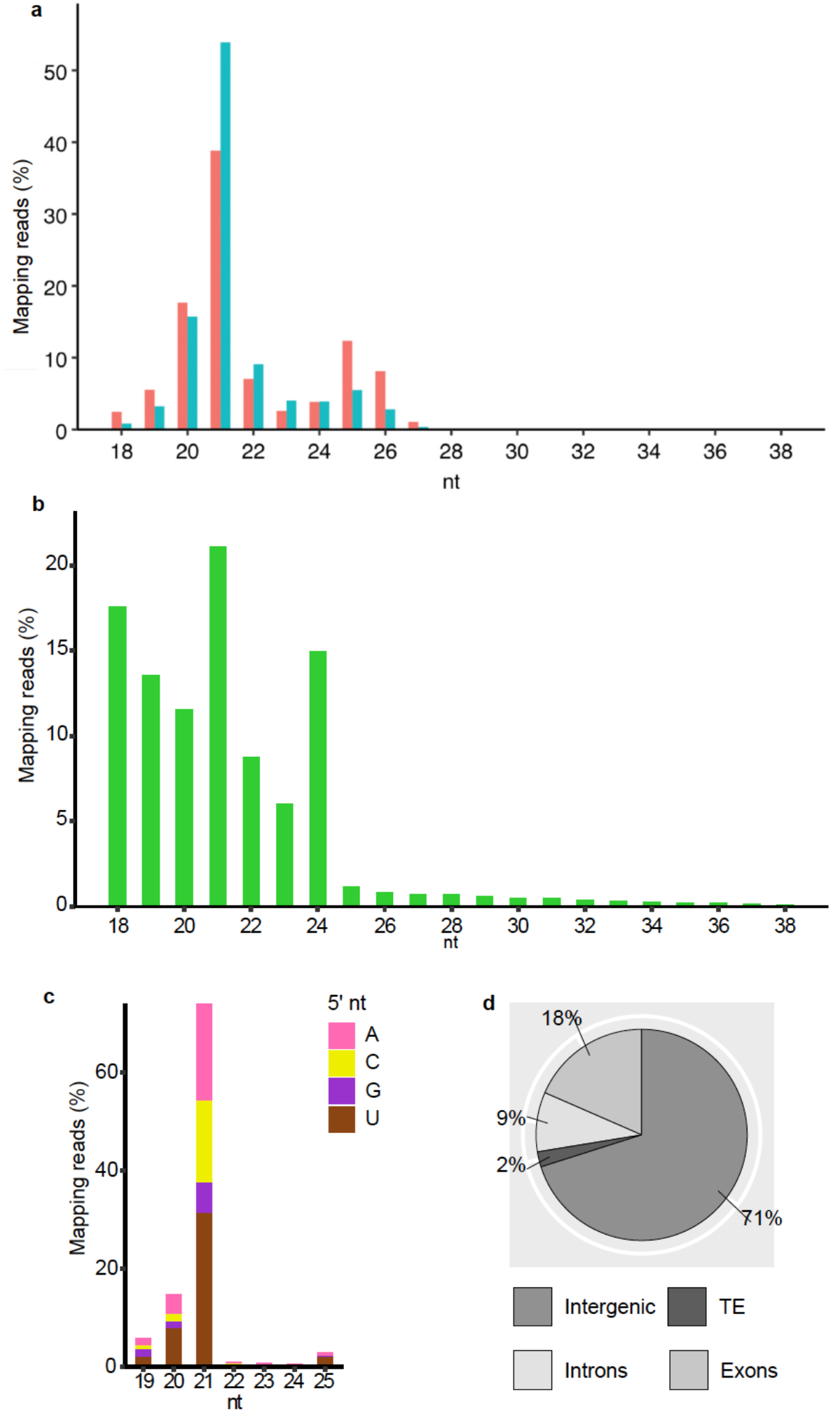
Characteristics of small RNAs derived from co-IP RNA samples of *P. infestans* (*Pi*) Ago1-GFP in mycelia and during infection of potato, *Solanum tuberosum* (*St*). **a,** *Pi*-sRNA in mycelia (blue, 15,863,929 processed reads) and during potato (cv. Bintje) infection (red, 5,412,731 processed reads). **b,** *St*-sRNA during potato infection (green, 1,432,780 processed reads). **c,** Identity of 5’ terminal nucleotide (nt) of *Pi*-sRNA mapping to potato genes, 5 days post infection. Percentages of *Pi*-sRNA per nt length, mapping to mRNAs with annotated gene functions, as visualized in Supplementary Fig. 4a. Distribution of 5’ U: 36% of 19 nt sRNAs, 54% of 20 nt sRNAs, 44% of 21 nt sRNAs and 77% of 25 nt sRNAs. **d,** *St*-sRNA mapping to the potato genome^16^. The target distribution is as following: 451 sRNA to transposable elements (TEs), where retrotransposons dominate (68% of all TEs), 13,558 sRNA to intergenic regions, 5338 sRNA to genes, divided on 3,573 sRNA to exons and 1,765 sRNA to introns. Duplicates were excluded in all datasets.

**Figure 2.**
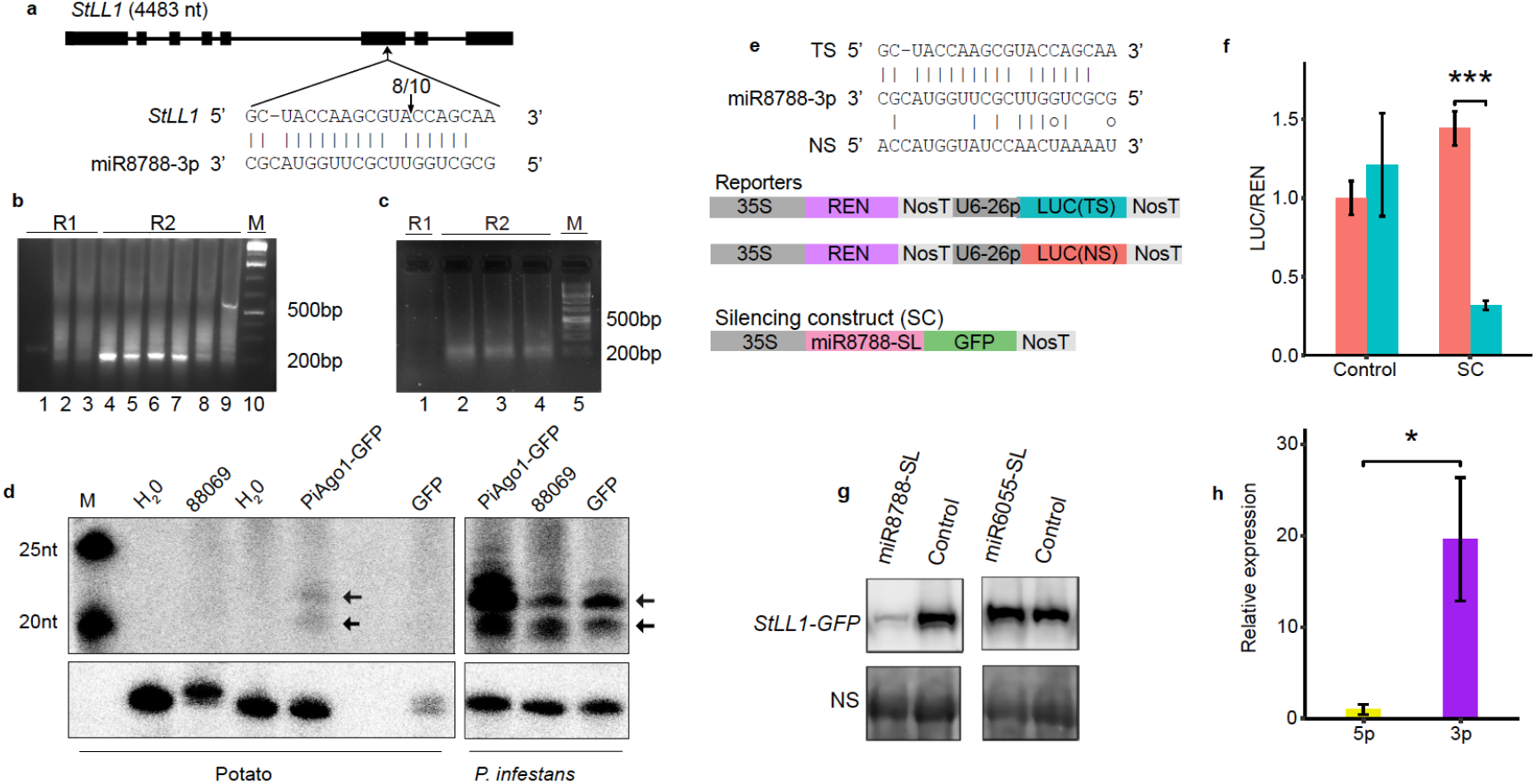
*StLL1* down-regulation is controlled by miR8788. **a,** Exons (filled box) and introns (black line) illustrating the lipase-like gene in potato (*StLL1*). Alignment of miR8788-3p with *StLL1* at the predicted binding site. The arrow with fraction above (8/10) indicates the cleavage site with the number of identical clones detected by 5’ RACE. **b,** 5’ RACE products of *StLL1* in *P. infestans*-infected potato (cv. Bintje). R1 (lane 1-3) achieved using GeneRacer™ 5’ primer and target mRNA 3’ primer. R2 (lane 4-9), miR8788-specific product generated with GeneRacer™ 5’ nested primer and target mRNA 3’ primer. **c,** 5’ RACE products of *StLL1* in water-inoculated potato (cv. Bintje). R1 (lane 1) achieved using GeneRacer™ 5’ primer and target mRNA 3’ primer. R2 (lane 2-4), products generated with GeneRacer™ 5’ nested primer and target mRNA 3’ primer. M = 1 kb Plus DNA. **d**, Northern blot analysis using a γ-^32^P labelled RNA probe for miR8788-5p. M = γ-^32^P labeled GeneRuler Ultra Low Range DNA Ladder. H20 = water inoculation. *P. infestans* infected potato leaves (with strains 88069, *pHAM34*:*PiAgo1-GFP* and *pHAM34:eGFP* (*GFP*). Samples (cv. Bintje) collected 5 dpi. miR8788-5p is indicated with black arrows. Lower panel = U6 snRNA from potato (loading control, probe cross-reacts with *P. infestans* U6). **e,** T-DNA constructs used in the luciferase reporter assay. 35S promoter (35S), REN luciferase (REN), U6 snRNA promoter (U6-26p), firefly luciferase (LUC), miR8788 target sequence in *StLL1* (TS), non-specific sequence (NS) and Nos 3’ terminator (NosT). **f,** Luciferase reporter assay in *N. benthamiana* samples, 3 days post Agro-infiltration. Agro-infiltration with GV3101 (Control) or silencing construct (SC). Reporters: *p35S:REN:LUC*(TS), blue, *p35S:REN:LUC*(NS), red. The quantified LUC normalized to REN activities are shown (LUC/REN). Error bars indicate mean ± standard error of the mean (*n* = 5, *df* = 19). *** = significant difference between the reporters during Agro-infiltration with SC (Student’s *t-*test: *P* < 0.001). **g,** Detection of StLL1-GFP protein co-infiltrated with miR8788-SL or with miR6055-SL (SL = stem loop). Control = Agro-infiltration of strain GV3101. NS= (non-specific) *N. benthamiana* Rubisco protein. Samples in the Western blot are probed with anti-GFP antibody. **h,** Relative transcript levels of miR8788-5p (yellow) and miR8788-3p (purple) in *N. benthamiana*, 3 days post miR8788-SL Agro-infiltration. Error bars indicate mean ± standard error of the mean (*n* = 4, *df* = 7). * = significant difference between transcript levels of miR8788-5p and miR8788-3p 3 days post Agro-infiltration (Student’s *t-*test: *P* < 0.05).

### mRNAs coding for resistance proteins are the dominating sRNA target

Concerning the origin of the different sRNAs loaded into PiAgo1, 71% of the *St*-sRNA derived from intergenic regions in the potato genome, followed by exons (18%) and introns (9%) (Fig.1d). Six *St*-miRNAs derived from potato protein-coding genes (4 exonic and 2 intronic) whereas the 27 additional *St*-miRNAs originated from intergenic regions.

We predicted mRNA targets and their annotated gene functions in potato for *Pi*-sRNAs, *St-* miRNAs and a remaining group of *St-*sRNAs having a greater coverage than 10 RPM. Genes coding for transporters, resistance proteins and kinases were top-three groups for *Pi*-sRNAs and *St*-sRNAs (Fig. S4). A bias towards the *R* gene category, 141 and 55, was detected in the two datasets. In the *St*-miRNA group, 21 miRNAs were found to target 107 *St-*genes whereof 34 were *R* genes (Fig. S4) followed by transcription factors and liguleless, the latter important for leaf development (Osmont *et al.*, 2003). In summary, as many as 206 *R* genes could be potentially down-regulated upon *P. infestans* infection either by the host or by pathogen-derived sRNAs loaded into PiAgo1 (Fig. S5a). In the *St*-sRNA and the *Pi*-sRNA datasets, 13 and 12 *R* genes, respectively, were annotated as late blight resistance genes, and three *Rpi-blb2-like* genes were targeted by both sources of sRNAs (Table S3). The genome of *P. infestans* is rich in transposable elements (TEs) (Haas *et al.,* 2009). Long terminal repeat (LTR) retrotransposons was earlier found as a major source of sRNAs in *P. infestans* mycelia (Vetukuri *et al.,* 2012). Here, during potato infection, 6% of the *Pi*-sRNAs targeting *R* genes originated from TEs, all derived from *Gypsy* LTRs. Relative predicted cleavage sites showed a bias towards the 5’ ends of the mRNAs for *R* genes in all three sRNA data sets, particularly pronounced in the two endogenous *St*-sRNA and *St*-miRNA subgroups (Fig. S5b). Among the 1,033 sRNAs from *P. infestans*, its single miRNA (miR8788) was found, being detected in the *pHAM34*:*PiAgo1-GFP* infected leaf sample, but not in the *pHAM34:eGFP* infected sample. None of the earlier predicted *Pi*-miRNAs (Cui *et al.*, 2014) were present in our reads. The miR8788-5p is located in exon 1 and the miR8788-3p is found in the short (57 nt) intron 1 of *PITG_10391*. Analysis of miR8788 (both 3p and 5p) revealed that its potential targets are a handful of genes in the *P. infestans* genome (Table S4) and a lipase-like gene (PGSC0003DMG400007679, henceforth *StLL1*) in the potato genome. Only miR8788-3p, but not miR8788-5p or any PiAgo1-bound miR8788-3p isomiRs, were predicted to target *StLL1*.

### miR8788 from *P. infestans* cleaves mRNA of potato lipase-like gene *StLL1*

To validate the predicted miR8788 cleavage site in *StLL1*, 5’ RACE was performed, using total RNA extracted from leaves 5 days post inoculation with *P. infestans*. The amplified PCR products, ranging from 100-200 bp, were cloned, sequenced and mapped to the potato genome (Hardigan *et al.*, 2016). Eight out of 10 clones terminated at the same position (Fig. 2a,b), supporting the miRNA-induced cleavage prediction. 5’ RACE on total RNA extracted from potato inoculated with water resulted in 40 identical clones, none of which indicated *StLL1* cleavage without *P. infestans* (Fig. 2c). The result was followed up by Northern hybridization, demonstrating the presence of miR8788-5p in *pHAM34*:*PiAgo1-GFP* infected potato (Fig. 2d). Involvement of any potential phasiRNA generated from the potato *R* genes targeting *StLL1* was also checked, but no such candidate was found. Next, the dual-luciferase reporter system was applied on Agro-infiltrated *Nicotiana benthamiana* leaf materials (Fig. 2e). This approach generated repression of the target mRNA sequence (Fig. 2f), also seen in Western blot analysis (Fig. 2g). Alternative miR8788 targets were searched for and three potential sequence candidates were analysed but no cleavage sites were detected in the predicted miRNA sites by 5’ RACE (Fig. S6).

### LL1 is prevalent among *Solanaceae* species

The StLL1 protein has a single trans-membrane domain and an alpha/beta hydrolase fold domain and is significantly suppressed in potato upon *P. infestans* inoculation (Fig. S7a,b). To determine StLL1 localization in *N. benthamiana* cells, a GFP-tag was fused to the C-terminus of StLL1 and the fusion protein was expressed under the control of the 35S promoter. After Agro-infiltration, StLL1-GFP was detected in the tonoplast (Fig. S7c-g), the membrane separating the vacuole from the cytoplasm. The tonoplast is an important subcellular compartment and point of contact with the specific intercellular infection structure called haustorium formed by oomycetes as invaginated plant host cells. In the Arabidopsis-*Hyaloperonospora arabidopsidis* interaction, the plant cellular content may undergo extensive rearrangements, including re-localisation of the tonoplast close to the haustorium under infection (Caillaud *et al.,* 2012). Much remains to be clarified on the haustorium-host molecular exchanges.

*StLL1* is member of a small family of two genes, located in different *Solanaceae* gene clades (Fig. 3; Fig. S8). A gene that appears fixed due to lack of positive selection for any amino acid of StLL1 using site models implemented in PAML (Yang, 2007). Neither was any positive selection found for the StLL1 branch. *StLL1* is also present in seven common potato cultivars, and the miR8788 target site was found in all genotypes (Fig. S9). Altogether, we propose that StLL1 is a conserved multifunctional protein with critical cellular functions in potato and other plant species, most likely associated with tonoplast-vacuole transport of critical compounds such as nutrients, ions and regulatory molecules, including sRNAs.

**Figure 3.**
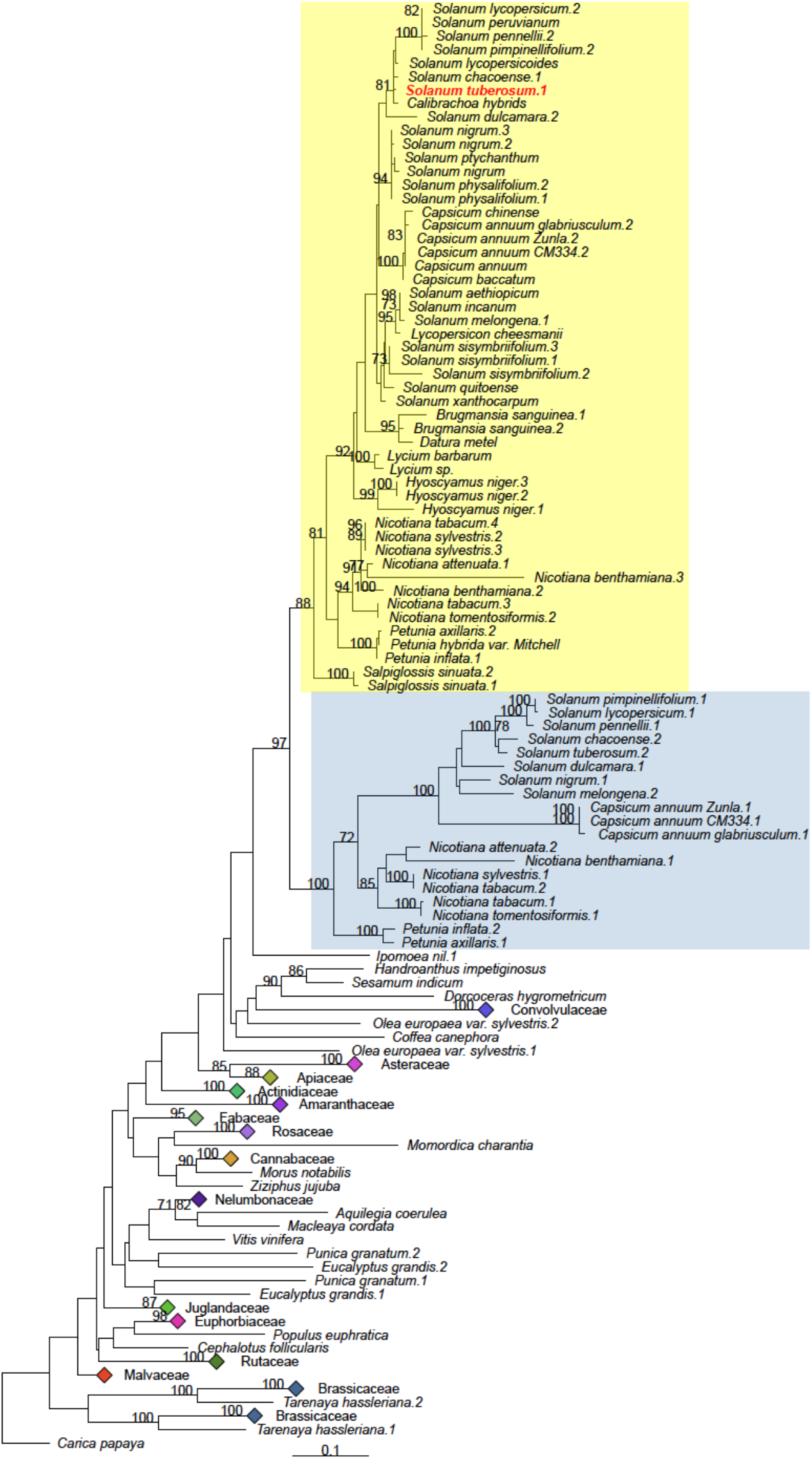
Unrooted maximum likelihood phylogeny of lipase-like (LL)-encoding genes. Bootstrap values > 70% are shown. Blue and yellow backgrounds highlight the Solanaceae subclades. Solanaceae sequences not found at NCBI were mined from https://solgenomics.net, *www.onekp.com,* www.ebi.ac.uk (PMID:26442032) and www.doi.org/10.5061/dryad.kc835.5. (RAxML, model JTT +Γ) was applied. Bar = number of substitutions per site. Bootstrap values > 70% are indicated. StLL1 (XP_006357616.1) is displayed in red.

### StLL1 is a critical defence component to *P. infestans*

Stable transgenic potato lines were produced using *Agrobacterium*-mediated transformation, first using the same StLL1 over-expression (OE) construct as in the transient *N. benthamiana* assay (Fig. 4a). These plants were used to determine the disease response to *P. infestans*. Inoculated leaves showed significantly smaller disease lesions and less growth of *P. infestans*, after 5 days (Fig. 4b,c). At the same time-point the *StLL1* transcript levels were about 10-fold lower compared to mock treatment (Fig. 4d). Although we cannot exclude the involvement of an endogenous potato factor in *StLL1* silencing, the data shows that StLL1 is critical for *P*. *infestans* defence. We also generated stable potato transgenic lines containing an artificial miRNA (*StamiRNA*) construct to induce silencing of the *StLL1* transcript (Fig. 4e). In contrast to our OE materials, *P. infestans* spread quickly in these transgenic lines (Fig. 4f). Already 2 days after inoculation, sporangiophores started to protrude from the leaves. After 3 days, many leaves were completely covered with mycelia, sporangiophores and sporangia. The DNA content of *P. infestans* was also significant higher in the *StamiRNA* plants when *StLL1* was knocked down compared to *StLL1* in wild-type potatoes (Fig. 4g). Similarly the *StLL1* transcript levels were significantly reduced in the *StamiRNA* plants (Fig. 4h). A miRNA target mimic approach was next applied to inhibit miR8788 activity in *P. infestans*. Six knock-down (KD) candidates were achieved, whereof KD1 was chosen and used for potato inoculations. We found that this *P. infestans* KD strain had reduced growth and lower miR8788 transcript levels on its host compared to the wild-type strain 88069 (Fig. 4i-k; Fig. S10), demonstrating the importance of miR8788 for the infection process.

**Figure 4.**
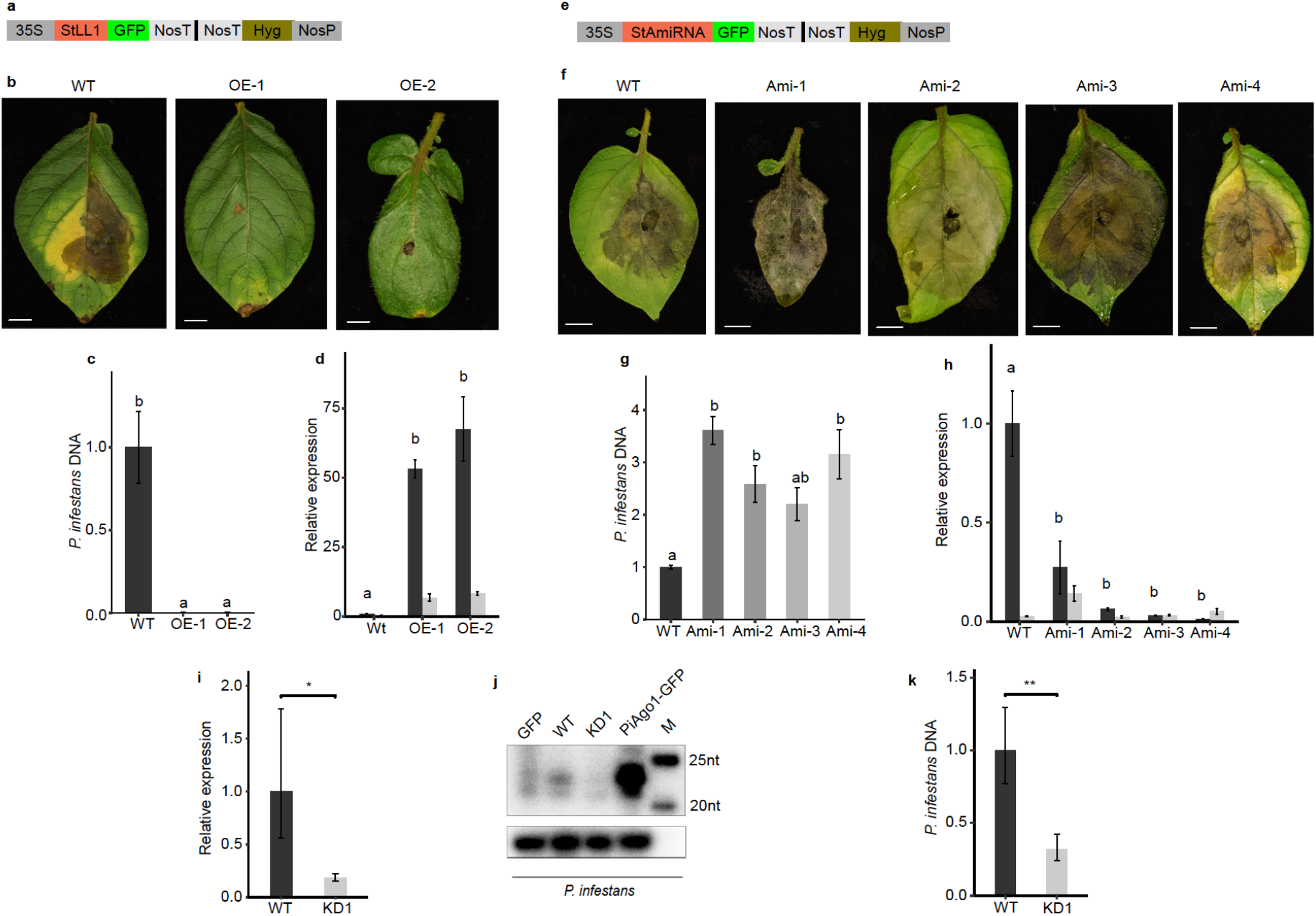
*StLL1* is essential for potato defence against *P. infestans*. **a,** Construct used to produce *StLL1* over-expression lines: 35S promoter, Green Fluorescent Protein (*GFP)*, Nos 3’ terminator (*NosT*), Hygromycin B resistance gene (*Hyg*), Nos promoter (*NosP*). **b,** Disease phenotype of wild-type (WT, cv. Sarpo Mira, strain 11388) and two transgenic over-expression lines containing the *p35S:StLL-GFP* construct (OE-1, OE-2), inoculated with *P. infestans*. Scale bar = 1 cm. **c,** Relative *P. infestans* DNA content in leaves from the over-expression potato lines in b. Error bars indicate mean ± standard error of the mean (*n* = 3, *df* = 8). Letters in the bar charts (a–b) represent significant differences (one-way ANOVA and Tukey’s HSD test: *P* < 0.001). **d,** Relative expression of *StLL1* in leaves from the over-expression lines in b, when inoculated with water (black) or *P. infestans* (grey). Error bars indicate mean ± standard error of the mean (*n* = 3, *df* = 17). Letters in the bar chart (a, b) represent significant differences (two-way ANOVA and Tukey’s HSD test: *P* < 0.001). **e,** Construct used to produce transgenic artificial miRNA lines: 35S promoter (*35S*), *GFP*, *StamiRNA*, Nos 3’ terminator (*Nos*T), Hygromycin B resistance gene (*Hyg*), Nos promoter (*NosP*). **f,** Disease phenotype of wild-type (WT, cv. Desirée, strain 88069) and four transgenic artificial miRNA potato lines (Ami-1, Ami-2, Ami-3, Ami-4) inoculated with *P. infestans*. Scale bar = 1 cm. **g,** Relative *P. infestans* DNA content in leaves from the artificial miRNA lines in f. Error bars indicate mean ± standard error of the mean (*n* = 3, *df* = 14). Letters in the bar chart (a, b) represent significant differences (one-way ANOVA and Tukey’s HSD test: *P* < 0.05). **h,** Relative expression of *StLL1* in leaves from the artificial miRNA potato lines in f, during inoculation with water (black) or *P. infestans* (grey). Error bars indicate mean ± standard error of the mean (*n* = 3, *df* = 19). Letters in the bar chart (a, b) represent significant differences (two-way ANOVA and Tukey’s HSD test: *P* < 0.001). **i,** Relative transcript levels of miR8788-3p in 88069 (WT) and miR8788 knock-down (KD1) mycelia. Error bars indicate mean ± standard error of the mean (*n* = 5, *df* = 9). * = significant difference between transcript levels of miR8788-3p between WT and KD1 (Student’s *t-*test: *P* < 0.05). **j,** Northern blot analysis using a γ-^32^P labeled RNA probe for miR8788-5p. *P. infestans* strains: *pHAM34:eGFP* (*GFP*), 88069 (WT), miR8788 knock-down (KD1), and *pHAM34*:*PiAgo1-GFP (PiAgo1-GFP)*. M = γ-^32^P labelled GeneRuler Ultra Low Range DNA. Lower panel = 5S rRNA from *P. infestans* (loading control). **k***, P. infestans* DNA content in potato leaves of cv. Bintje infected with *P. infestans* strain 88069 (WT, black) and miR8788 knock-down (KD1). Error bars indicate mean ± standard error of the mean (*n* = 7, *df* = 13). ** = significant difference in *P. infestans* DNA content between WT and KD1 (Student’s *t-*test: *P* < 0.01). Leaves were sampled 5 dpi.

## Discussion

*P. infestans* miR8788 is located in the *PITG_10391* sequence, encoding a protein with unknown function. *PITG_10391* is a unique single gene in the genome of *P. infestans* located on supercontig 1.18, a genomic region with low frequency of transposons. Besides targeting the mRNA of *StLL1,* miR8788 is also predicted to target mRNAs in the *P. infestans* genome, particularly for the amino acid/auxin permase (AAAP) family. AAAPs are found in almost all eukaryotes. These proteins contribute to various stress responses and long distance amino acid transport, the latter which can be mediated across the cellular membrane (Wipf *et al.,* 2002; Tegeder, 2012). During tuber infection, 54 out of 57 *AAAP* genes predicted in the *P. infestans* genome were active particularly PITG_20230 and PITG_12808 (Ah-Fong *et al.,* 2017). How *AAAP* genes are regulated in eukaryotes remains to be clarified.

*P. infestans* is known to encode hundreds of effector proteins, those targeted to the host apoplasm like cell wall degrading enzymes and plant protease inhibitors or those that translocate into plant cells (Kamoun, 2008). The latter cytoplasmic effector category is divided into two main groups of Crinklers (CRNs) and those with the conserved Arg–any amino acid–Leu–Arg (RXLR) peptide motif (Haas *et al.*, 2009). A handful of RXLR effectors were studied more closely and found localised to a variety of host plant cell compartments such as the cytoplasm and nucleoplasm (Bos *et al.*, 2010), the endoplasmic reticulum (McLellan *et al.*, 2013) and many other places (Wang *et al.*, 2019). This diversity of target sites highlights the possible power of concerted action during infection of *P. infestans*. However, there is a need to move from studies in *N. benthamiana* to potato harbouring various resistance genes to clarify the importance of all targets described.

Mechanisms of effector delivery to plant cells are not completely understood despite evidence that an RXLR effector can be secreted to the inside of the plant cell (Whisson *et al.*, 2007). By using confocal microscopic approaches, two effectors (one cytoplasmic and the other apoplastic) both were enriched in the haustoria followed by secretion to *N. benthamiana* cells (Wang *et al.*, 2017). The apoplastic EPIC1 effector used the Golgi-mediated secretion pathway whereas the cytoplasmic Pi04314 used an alternative, still unknown pathway. Other studies have also reported that cytoplasmic effectors accumulate in the haustoria (Gilroy *et al.*, 2011; Liu *et al.*, 2014; Wang *et al.*, 2018, 2019a), which presently is believed to be the major site for effector secretion in *P. infestans*. Transport of the PiAgo1-GFP protein into plant cells has yet not been possible to clearly demonstrate by microscopic analysis. Similarly, the function of StLL1 is not known. The tonoplast surrounds the vacuole in plant cells and contains numerous transporters and function in vesicle trafficking (Wolfenstetter *et al.*, 2012). Whether or not the tonoplast plays a role in the sRNA trafficking between *P. infestans* and the host plant and the involvement of StLL1 in this process are intriguing future aspects on this study. Knowledge on sRNA translocation events is emerging from several plant-pathogen systems (Hudzik *et al.*, 2020). Among *Phytophthora* species, most information derives from *P. sojae* (Qutob *et al.,* 2013; Wang *et al.,* 2019b). Recently it was shown that in the Arabidopsis -*P. capsici* interaction, secondary sRNAs from the host were induced upon infection and targeted pathogen genes to promote defense. However in parallel, a known effector (PSR2) suppressed this production of secondary sRNAs and the processes resulted in enhances susceptibility (Hou *et al.,* 2019). Phytophthora suppressors of RNA silencing (PSRs) were originally found in *P. sojae* and are common among nine oomycete species according to phylogenetic analysis (Xiong *et al.*, 2014). *P. infestans* lags behind, much due to its large and complex genome, in combination with the demanding molecular toolbox presently available. Nevertheless, in light of its importance as plant pathogen, it is worth noting that when searching for information on *PITG_10391*, we found it being present with intact pre-miR8788 sequence in raw reads of European herbarium materials collected 1846 (strain KM177513) and 1877 (strain M-0182896) (Yoshida *et al.*, 2013). *PITG_10391* is still present in European *P. infestans* strains (Fig. S11), demonstrating its importance for this pathogen.

## Acknowledgements

The authors want to thank Dr. German Martinez Arias for valuable suggestions on the work and comments on the manuscript. The authors want to acknowledge, the *Solanum lycopersicoides* genome consortium for sequence information, and the support of the National Genomics Infrastructure (NGI)/Uppsala Genome Center and UPPMAX for providing assistance in massive parallel sequencing and computational infrastructure. This study was supported by the Swedish Research Council VR (2015-04259), Helge Ax:son Foundation and the Swedish University of Agricultural Sciences. Work performed at NGI /Uppsala Genome Center has been funded by RFI/ VR and Science for Life Laboratory, Sweden.

## Author contributions

X.H., A.Å., K.P.H., Z.L. and F.D. performed experiments and analysed data; X.H., A.Å., K.P.H., Z.L. and C.D. designed experiments; C.D., K.P.H. and A.Å. wrote the manuscript.

## Supporting information

Additional Supporting Information may be found online in the Supporting Information section at the end of the article.

The following Supporting Information is available for this article:

**Fig. S1** Relative *P. infestans* DNA content in infected potato.

**Fig. S2** Pipeline for the bioinformatics analysis.

**Fig. S3** Individual replicates of small RNAs derived from co-IP-enriched RNA samples.

**Fig. S4** Predicted sRNA target mRNAs in potato.

**Fig. S5** sRNA subgroups targeting resistance genes in the potato genome.

**Fig. S6** *PITG_10391* and miR8788 potato target candidate assay.

**Fig. S7** StLL1 analysis.

**Fig. S8** Phylogenetic tree of the lipase-like (LL) encoding genes.

**Fig. S9** Multiple sequence alignment of *StLL1* CDS in different potato cultivars.

**Fig. S10** Relative transcript levels of miR8788-5p in 88069 (WT) and miR8788 knock-down (KD1).

**Fig. S11** Genomic sequence comparisons of miR8788.

**Table S1** Oligo sequences for RACE, constructs and Northern blot.

**Table S2** qRT-PCR primer sequences

**Table S3** sRNAs predicted to target the same resistance genes in the potato genome

**Table S4.** Predicted miR8788 target transcripts in the *P. infestans* genome.

