## Supplementary material for "*Phytophthora infestans* Ago1-bound miRNA promotes potato late blight disease": Suppl. figures and tables

### New Phytologist Supporting Information Figs S1-S11 and Tables S1-S4

The following Supporting Information is available for this article:

**Fig. S1** Relative *P. infestans* DNA content in infected potato.
**Fig. S2** Pipeline for the bioinformatics analysis.
**Fig. S3** Individual replicates of small RNAs derived from co-IP-enriched RNA samples.
**Fig. S4** Predicted sRNA target mRNAs in potato.
**Fig. S5** sRNA subgroups targeting resistance genes in the potato genome.
**Fig. S6** *PITG_10391* and miR8788 potato target candidate assay.
**Fig. S7** StLL1 analysis.
**Fig. S8** Phylogenetic tree of the lipase-like (LL) encoding genes.
**Fig. S9** Multiple sequence alignment of *StLL1* CDS in different potato cultivars.
**Fig. S10** Relative transcript levels of miR8788-5p in 88069 (WT) and miR8788 knock-down (KD1).
**Fig. S11** Genomic sequence comparisons of miR8788.
**Table S1** Oligo sequences for RACE, constructs and Northern blot.
**Table S2.** qRT-PCR primer sequences.
**Table S3.** sRNAs predicted to target resistance genes in the potato genome.
**Table S4.** Predicted miR8788 target transcripts in the *P. infestans* genome.

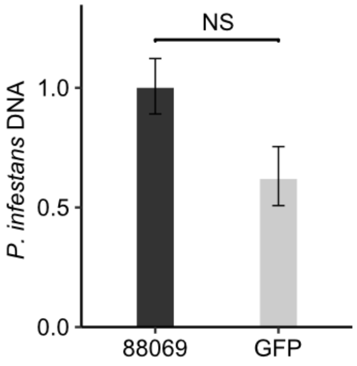


**Fig. S1** Relative *P. infestans* DNA content in potato cv. Bintje leaves infected with *P. infestans* strain 88069 (black) or *pHAM34:eGFP* (GFP) (grey). Error bars indicate mean ± standard error of the mean (n = 7). NS = no significant difference between the 88069 and GFP strains.

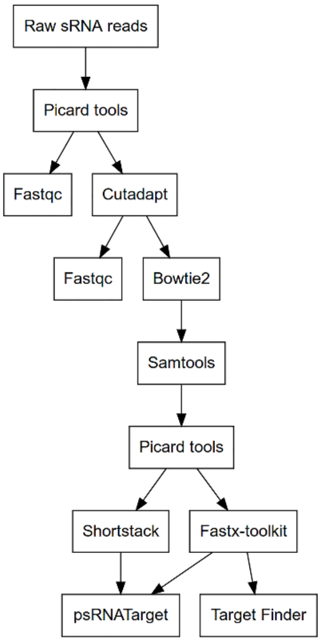


**Fig. S2 Pipeline for the bioinformatics analysis of the Co-IP enriched RNA derived from potato leaves inoculated with GFP-tagged *P. infestans*, sequenced on the Ion Proton platform.** Four raw data sets were analysed: transgenic *P. infestans* mycelia containing *pHAM34:PiAgo1-GFP* or *pHAM34:eGFP*, and potato leaves (cv. Bintje) inoculated with the same strains. The function SamToFastq, included in the Picard tool package, was used to convert the original BAM format to FASTQ format, on which a quality control was performed, using FastQC. Cutadapt was used for trimming with a quality cutoff of 20 using 18 nt and 38 nt as minimum and maximum read lengths. Bowtie2 in combination with SAMtools view were used for tRNA and rRNA trimming, allowing 0 mismatches per seed against each filtered dataset. The same tools were used to separate potato and *P. infestans* sRNA by first mapping the sRNA to each genome (Haas *et al*., 2009; Hardigan *et al*., 2016). Next, the *St*-sRNA pool was trimmed against the *P. infestans* genome and the *Pi*-sRNA pool was trimmed against the potato genome to exclude potential false positives. This procedure split the leaf dataset into two. Altogether, six sRNA datasets (*Pi*-sRNA from mycelia of *pHAM34:PiAgo1-GFP* and *pHAM34:eGFP*, *Pi*-sRNA from *pHAM34:PiAgo1-GFP* and *pHAM34:eGFP* infected leaves and *St*- sRNA from *pHAM34:PiAgo1-GFP* and *pHAM34:eGFP* infected leaves) were generated. SamToFastq converted the files to FASTQ format. ShortStack was used to predict miRNA and summarize sRNA, using strand cutoff 0.5, fold size 1000, dicermin 18 and dicermax 38. Fastx_collapser in the FASTX-toolkit counted the number of sRNA. psRNATarget was implemented to predict mRNA targets of sRNA and miRNA against the *Solanum tuberosum* genome transcript (JGI genomic project, Phytozome 12, 448_v4.03) with default settings, except the expectation value was 3. When running miR8788 the expectation value was 5. TargetFinder predicted mRNA targets of sRNA and miRNA in the genome of *P. infestans*.

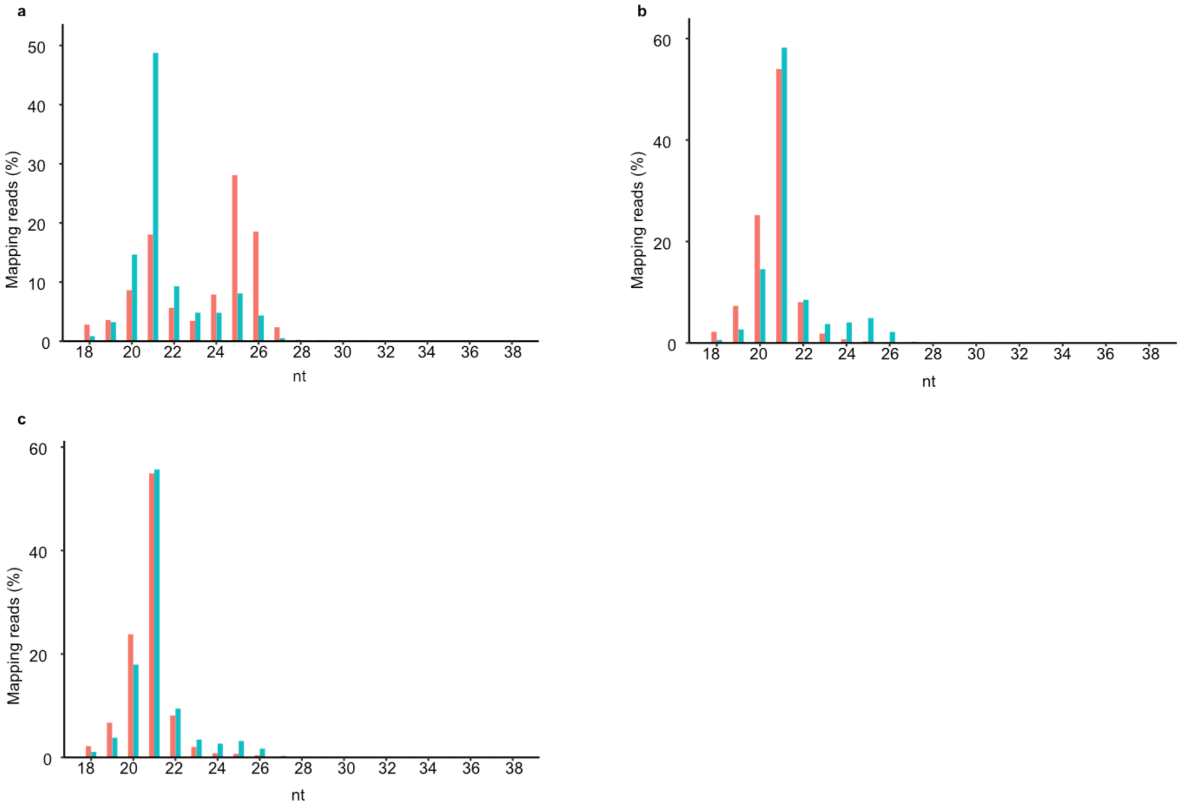


**Fig. S3 Individual replicates of small RNAs derived from co-IP-enriched RNA samples of PiAgo1-GFP mycelia and infected potato leaves as shown in Fig. 1a. a,** First replicate of *Pi*-sRNA in mycelia (blue, 5,241,112 processed reads), and during potato infection (red, 610,713 processed reads). **b,** Second replicate of *Pi*-sRNA in mycelia (blue, 6,110,909 processed reads), and during potato infection (red, 2,163,509 processed reads). **c,** Third replicate of *Pi*-sRNA in mycelia (blue, 4,511,908 processed reads), and during potato infection (red, 2,638,509 processed reads).

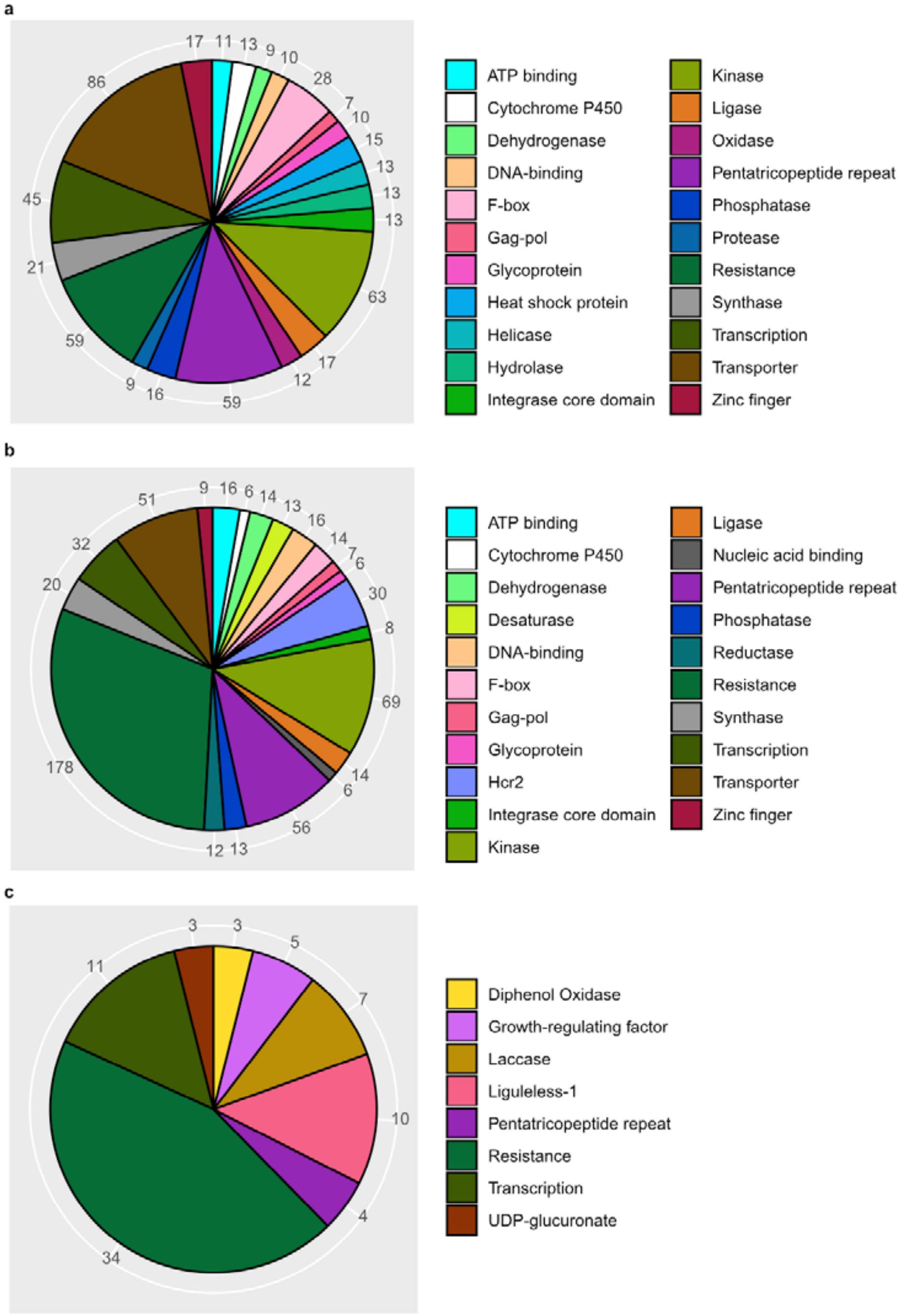


**Fig. S4 Predicted sRNA target mRNAs in potato organized according to annotated gene functional categories in the potato genome. a,** *Pi*-sRNA (1033) targeted genes in potato organized according to annotated gene functional categories in the potato genome. Each potato functional group contains > 6 similar genes. Out of 1,329 genes, 298 genes with unknown functions and 485 genes not fitting into any of these groups were removed. Gene numbers for each category are indicated at the fringe. **b,** *St*-sRNA (392) targeted genes in potato, excluding the *St*-miRNAs targeted genes in c. Each potato functional group contains > 6 similar genes. Out of 1,235 genes, 228 genes with unknown functions and 417 genes not fitting into any of these groups were removed. Gene numbers for each category are indicated at the fringe. **c,** *St*-miRNA (21) targeted genes in potato. Each potato functional group contains > 2 similar genes. Out of 107 genes, 8 genes with unknown functions and 22 genes not fitting into any of these groups were removed.

a b

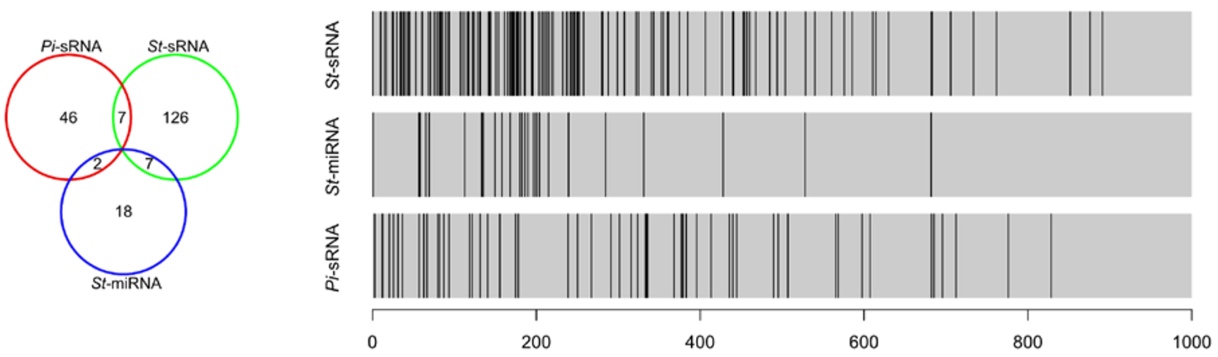


**Fig. S5 sRNA subgroups targeting resistance genes in the potato genome a,** Distribution of resistance gene mRNA, targeted by sRNAs, displayed in Supplementary Fig. 2**. b,** Relative target site positions in the *R* gene mRNA displayed in a.

S5

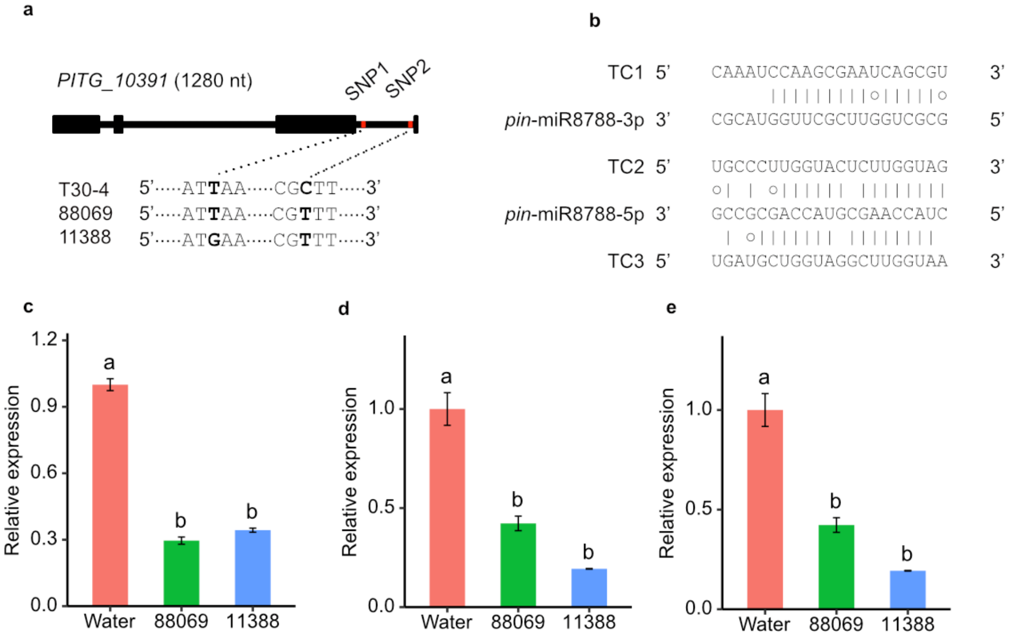


**Fig. S6 *PITG_10391* and miR8788 potato target candidate assay. a,** Comparison of SNPs in *PITG_10391* found in *P. infestans* strains 88069 and 11388 in relation to the reference strain (T30-4). Filled boxes = exons. **b,** Alternative gene target candidates (TC) of miR8788 in the potato genome. miR8788-3p was predicted to target TC1. miR8788-5p was predicted to target TC2 and TC3. TC1= PGSC0003DMT400056685, TC2= PGSC0003DMG400012409, TC3= PGSC0003DMG400018413. No cleavage sites were detected by 5' RACE. **c,** Relative transcript levels of TC1 in cv. Sarpo Mira inoculated with water, 88069 and 11388. Error bars indicate mean ± standard error of the mean ($n$ = 3, $df$ = 8). Letters in the bar chart (a, b) represent significant differences (one-way ANOVA and Tukey's HSD test: $P < 0.001$). **d,** Relative transcript levels of TC2 in cv. Sarpo Mira inoculated with water, 88069 and 11388. Error bars indicate mean ± standard error of the mean ($n$ = 3, $df$ = 8). Letters in the bar chart (a, b) represent significant differences (one-way ANOVA and Tukey's HSD test: $P < 0.001$). **e,** Relative transcript levels of TC3 in cv. Sarpo Mira inoculated with water, 88069 and 11388. Error bars indicate mean ± standard error of the mean ($n$ = 3, $df$ = 8). Letters in the bar chart (a–c) represent significant differences (one-way ANOVA and Tukey's HSD test: $P < 0.01$).

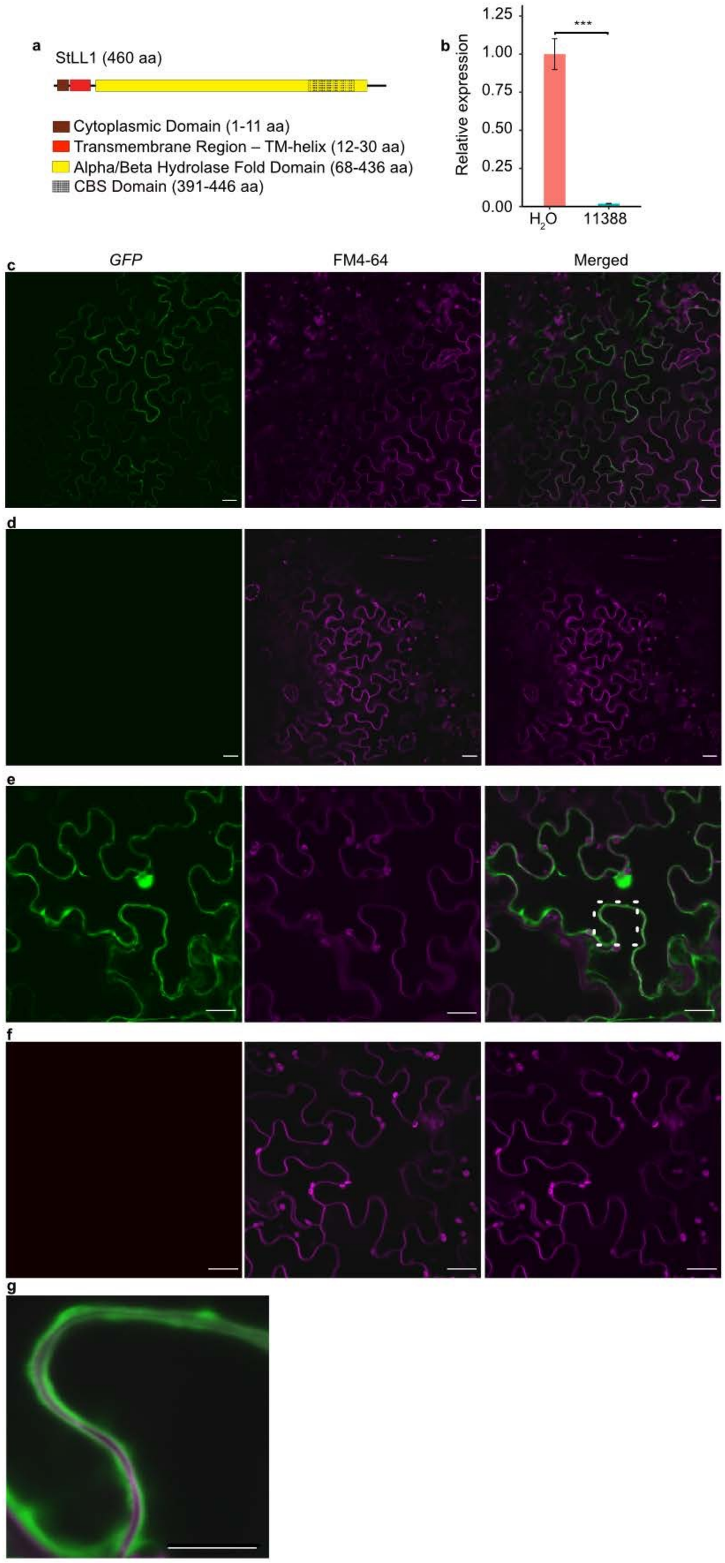


**Fig. S7 StLL1 analysis. a,** Illustration of predicted StLL1 domains using Interpro (http://www.ebi.ac.uk/interpro/). TMHMM Server version 2.0 predicted one transmembrane helix between amino acid residues 12 and 34 with greater than 90% certainty. (http://www.cbs.dtu.dk/services/TMHMM/). The amino acid residues downstream of amino acid number 34 are predicted to be extravacuolar. **b,** *StLL1* transcript analysis in potato cv. Sarpo Mira inoculated with *P. infestans* strains 11388 and $H_2O$, 5 dpi. Error bars indicate mean ± standard error of the mean (*n* = 3, *df* = 5). *** = significant differences (Student's t-test: *P* < 0.0001). **c,** Subcellular localization of the GFP-tagged StLL1 protein co-infiltrated with p19 in *N. benthamiana.* **d,** Agro-infiltration of p19 alone in *N. benthamiana*. **e,** Subcellular localization of the GFP-tagged StLL1 protein co-infiltrated with p19 in *N. benthamiana*. White square indicates area shown in g. **f,** Agro-infiltration of p19 alone in *N. benthamiana*. Scale bar in c-f = 20 µm. ***g,*** Area zoomed in from e. Scale bar = 10 µm. The plasma membrane was stained with FM4-64FX (FM4-64). Chlorophyll shows autofluorescence in the FM4-64 channel.

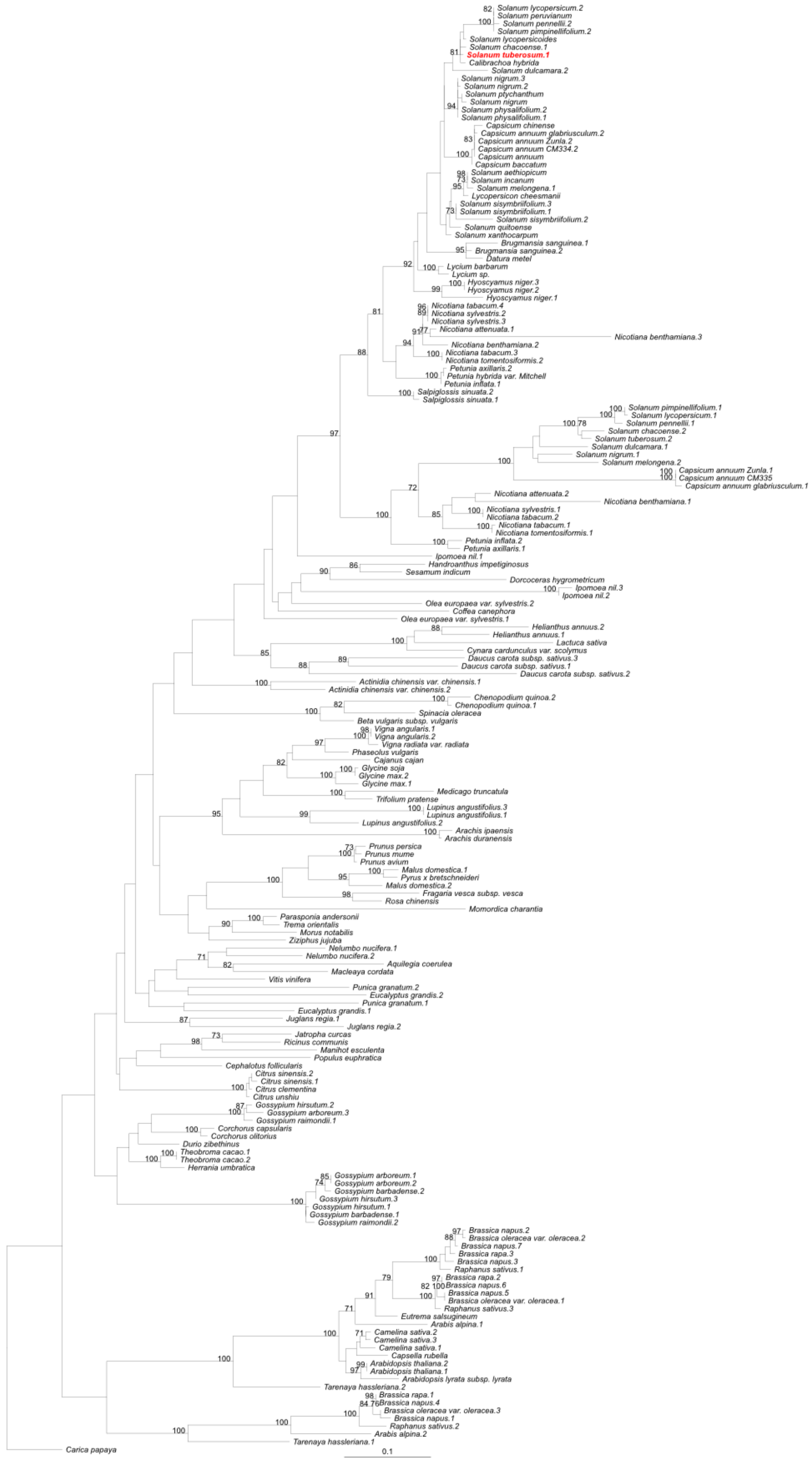

**Fig. S8 Phylogenetic tree of the unrooted maximum likelihood phylogeny (RAxML, model JTT +Γ) of the lipase-like (LL) encoding genes**. Bootstrap values > 70% are indicated. StLL1 (XP_006357616.1) is shown in red. Bar = number of substitutions per site. Pairwise alignment of StLL1 and StLL2 sequences, respectively, generated the following identity scores (%) with *A. thaliana* AtLL1 (74.4, 72.4) and AtLL2 (68.0, 73.9), and StLL1 vs. StLL2 = 81.5 %.

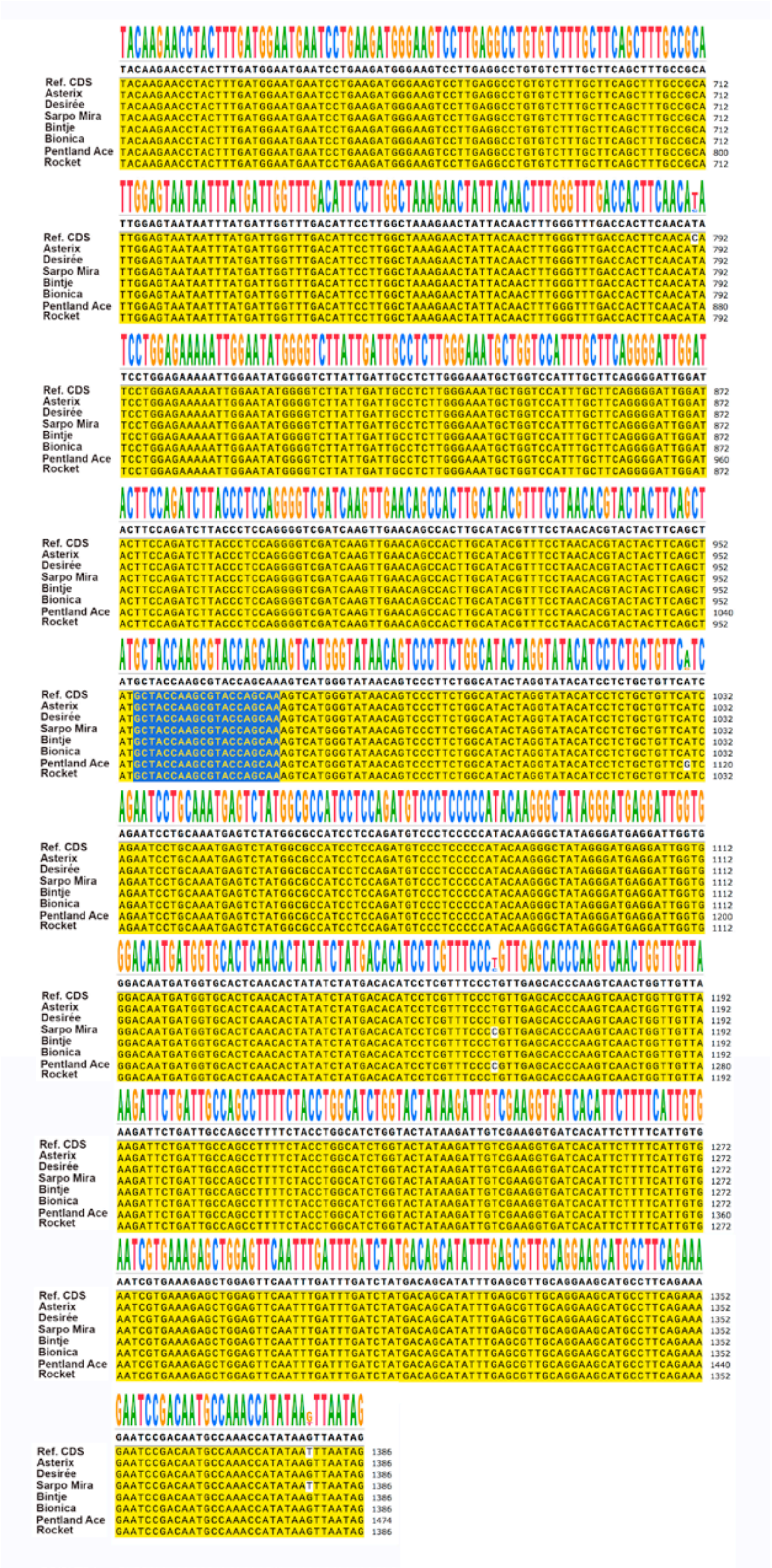

**Fig. S9 Multiple sequence alignment of *StLL1* CDS in different potato cultivars.** Around 320 bp upstream and downstream of the miR8788 target site were used for multiple sequence alignment. The miR8788 target site in *StLL1* is marked in blue (GCTACCAAGCGTACCAGCAA). The *StLL1* CDS was cloned from seven potato cultivars (Asterix, Desirée, Sarpo Mira, Bintje, Bionica, Pentland Ace and Rocket). Bases divergent from reference CDS in PGSC v4.04 are highlighted in white.

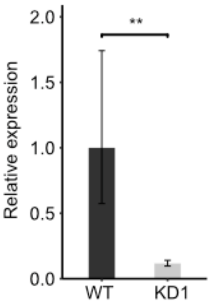


**Fig. S10 Relative transcript levels of miR8788-5p in 88069 (WT) and miR8788 knock-down (KD1) in mycelia**. Error bars indicate mean ± standard error of the mean (*n* = 5, *df* = 9). ** = significant difference between transcript levels between WT and KD1 (Student's t-test: $P < 0.01$).

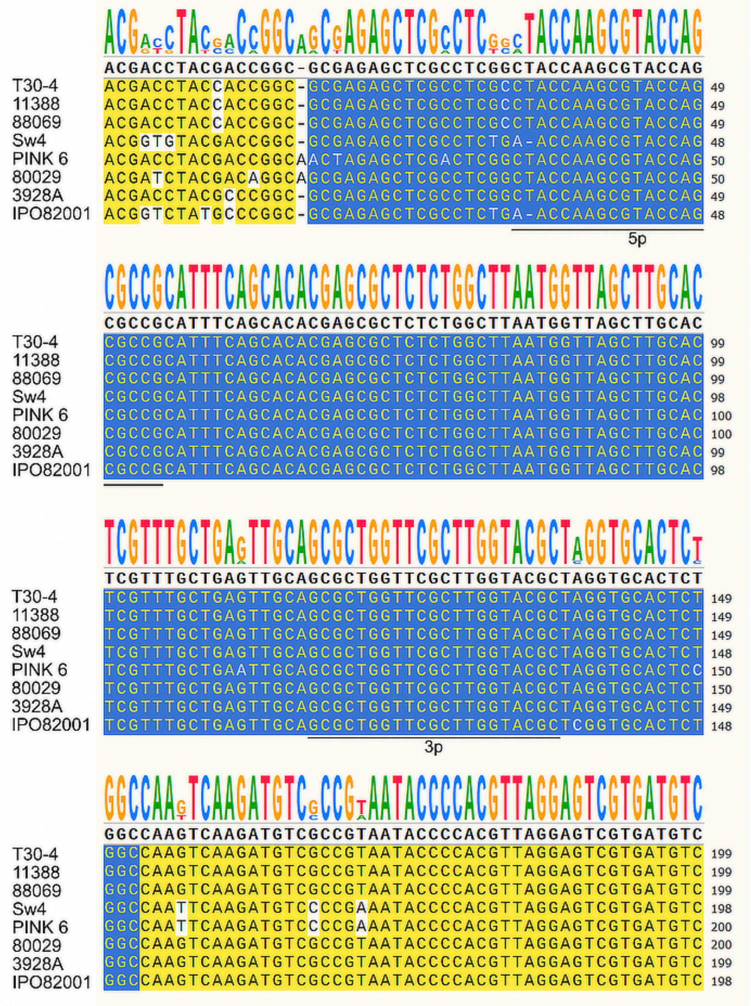

**Fig. S11 Genomic sequence comparisons of miR8788.** Multiple sequence alignment of the *PITG_10391* genetic sequence in eight *P. infestans* strains. The conserved miR8788 stem-loop is shown in blue. miR8788-5p and miR8788- 3p are marked by black lines. Nucleotide differences compared to the reference sequence in T30-4 are shown in white.

**Table S1. Oligo sequences for RACE, constructs and Northern blot**

| Name | Template | Sequence 5'-3' | Assays |
|---|---|---|---|
| TC1 -500BP-R | *TC1* | 5'-TTTGCTGCTACTGACTCTAAGGTT -3' | RACE |
| TC1 -200BP-R | | 5'- GATCTAAGTAGTTGAAAGCTATCGTTG -3' | |
| TC2 -500BP-R | *TC2* | 5'-CACCACCCTCTCTAGATCCAA-3' | |
| TC2 -200BP-R | | 5'-TTTCATAAATTCCAAGGGCAAG-3' | |
| GeneRacer™ RNA Oligo | GeneRacer™ RNA Oligo | 5'-CGACTGGAGCACGAGGACACTGACATGGACTGAAGGAGTAGAAA-3' | |
| GeneRacer™ 5′ Primer | | 5'-CGACTGGAGCACGAGGACACTGA-3' | |
| GeneRacer™ 5′ Nested Primer | | 5'-GGACACTGACATGGACTGAAGGAGTA-3' | |
| StLL1 -500BP-R1 | *StLL1* | 5'-GTGTTTGTAGCGCGCATTAG-3' | |
| StLL1 -200BP-R2 | | 5'-AACCAGTTGACTTGGGTGCT-3' | |
| StLL1-F | *StLL1* | 5'-CACCATGATAGTTAGGTGGTGGATAACAGC-3' | Constructs |
| StLL1-R1 | | 5'-AATTATATGGTTTGGCATTGTCG-3' | |
| StLL1-R2(TGA) | | 5'-TTAAATTATATGGTTTGGCATTGTC-3' | |
| miR8788-SL-F | miR8788-SL | 5'-CACCTACGATGTCCACACTCTCGAT-3' | |
| miR8788-SL-R | | 5'-CTCATGCTATCGCACGACAT-3' | |
| PITG_10391- F | *PITG_10391*- Genomic | 5'-CACCATGGCCAGCCAGAATCCGTCCAC-3' | |
| PITG_10391- R | | 5'-TCACGTTGTTGCTGAAGTTG-3' | |
| pU6-F | U6-Promoter | 5'-GGGGTACCAAACTCGACTTGCCTTCC-3' | |
| pU6-R | | 5'-ATAAGAATGCGGCCGCCAATCACTACTTCGACTCTAGCTG-3' | |
| LUC(TS)-F | *LUC*(TS) | 5'-ATAAGAATGCGGCCGCATGGAAGACGCCAAAAACAT-3' | |
| LUC(TS)-R | | 5'-GCTCTAGATTATTGCTGGTACGCTTGGTAGCCACGGCGATCTTTCC-3' | |
| LUC(NS)-F | *LUC*(NS) | 5'-ATAAGAATGCGGCCGCATGGAAGACGCCAAAAACAT-3' | |
| LUC(NS)-R | | 5'-GCTCTAGATTATTGCGCCGATTTGTGGTAGCCACGGCGATCTTTCC-3' | |
| 35S-F | 35S-promoter | 5'- GGGGTACCGTCTCAGAAGACCAAAGGGCTA-3' | |
| 35S -R | | 5'-ATAAGAATGCGGCCGCTAGAGCTCTTATACTCGAGCGTGTC-3' | |
| Ami-StLL1-F | *StLL1* | 5'-TTTTGGATATCACAGATGCCC-3' | |
| Ami-StLL1-R | | 5'-GGGCATCTGTGATATCCAAAA-3' | |
| Ami-StLL1*-F | | 5'-GAGCATCCGTGAAATCCAAAT-3' | |
| Ami-StLL1*-R | | 5'-ATTTGGATTTCACGGATGCTC-3' | |
| Ami-StLL2-F | | 5'-TAACGCCTTATCAGGTAGCAT-3' | |
| Ami-StLL2-R | | 5'-ATGCTACCTGATAAGGCGTTA-3' | |
| Ami-StLL2*-F | | 5'-ACGCTACATGATTAGGCGTTT-3' | |
| Ami-StLL2*-R | | 5'-AAACGCCTAATCATGTAGCGT-3' | |
| A Oligo | | 5'-CACCCTGCAAGGCGATTAAGTTGGGTAAC-3' | |

| | | | |
|---|---|---|---|
| B Oligo<br><br>GFP-F<br><br>MIM8788a-R<br><br><br>MIM8788b-R | *P. infestans* miRNA mimic | 5’- GCGGATAACAATTTCACACAGGAAACAG-3’<br><br>5’-CGGAATTCATGGTGAGCAAGGGCGAGGAGCTGTT-3’<br><br>5’-AAGGAAAAAAGCGGCCGCGCGCTGGTGTCTATGGTACGCCTT GTACAGCTCGTCCATGC-3’<br><br>5’-AAGGAAAAAAGCGGCCGCGCGCTGGTTTCTATGGTACGCCTTG TACAGCTCGTCCATGC-3’ | |
| miR8788-5p<br><br>StU6<br><br>Pi 5S rRNA | miR8788<br><br>U6 snRNA<br><br>5S rRNA | 5’-GCGGCGCUGGUACGCUUGGUAG-3’<br><br>5’-GCTAATCTTCTCTGTATCGTTCC-3’<br><br>5’-GCTTAACTTCACAGAGCAGAC-3’ | Northern blot |

**Table S2. qRT-PCR primer sequences**

| Name | Template | Sequence 5'-3' | Assays |
|---|---|---|---|
| StLL1-Q-F | *StLL1* | 5'-GCCACTTGCATACGTTTCCT-3' | qRT-PCR |
| StLL1-Q-R | | 5'-CCACCAATCCTCATCCCTAT-3' | |
| StACT-Q-F | *StACT* | 5'-GCCTCCTGAACGGAAGTACA -3' | |
| StACT-Q-R | | 5'-AATGGAAGGACCGGATTCAT-3' | |
| miR8788-5′ -RT | miR8788- 5′ | 5'-GTCGTATCCAGTGCAGGGTCCGAGGTATTCGCACTGGATACG ACCGGCGC- 3' | |
| pin-5′-Forward Primer | | 5'-TCGCGCTACCAAGCGTACCA- 3' | |
| Reverse Primer | | 5'-GTGCAGGGTCCGAGGT- 3' | |
| miR8788-3′ -RT | miR8788- 3′ | 5'-GTCGTATCCAGTGCAGGGTCCGAGGTATTCGCACTGGATACG ACGCGTAC - 3' | |
| pin-3′-Forward Primer | | 5'- GCTTCGCGCTGGTTCGCTTG - 3' | |
| Pin-ACT-Q-F | *Pin-ACT* | 5'- CATCAAGGAGAAGCTGACGTACA - 3' | |
| Pin-ACT-Q-R | | 5'- GCAGCTCGTAGCTCTTCTCCA - 3' | |
| Pin-qRT-PCR | *Pi*O8 | F CAATTCGCCACCTTCTTCGA | |
| Pin-qRT-PCR | | R GCCTTCCTGCCCTCAAGAAC | |
| StTC1-Q-F | *TC1* | 5'- CGCGAAAAACTCAACGATAG -3' | |
| StTC1-Q-R | | 5'- ATCGCATCGGATTTCATCAT -3' | |
| StTC2-Q-F | *TC2* | 5'- TAGCAAGGATCGGAGTTGTTG -3' | |
| StTC2-Q-R | | 5'- CTTCACAGCATCCACACTCG-3' | |
| StTC3-Q-F | *TC3* | 5'- TGGGGATAAGGGTTTGGTTT -3' | |
| StTC3-Q-R | | 5'- TCTCCTGTACACCCGAATCC-3' | |
| NtUbi-Q-F | *NtUbi* | 5'- TCCTGATGGGCAAGTGATTAC -3' | |
| NtUbi-Q-R | | 5'- TTGTATGTGGTCTCGTGGATTC-3' | |
| NtACT-Q-F | *NtACT* | 5'- AATGTGAAAGCCAAGATCCAAG-3' | |
| NtACT-Q-R | | 5'-CGGAGGCGGAGCACGAGATGAA-3' | |

**Table S3. sRNAs predicted to target resistance (*R*) genes in the potato genome distributed as illustrated in Fig. S4a.**

**a,** Seven predicted *R* genes shared by the *St*-sRNA and *Pi*-sRNA pools

| **Target gene ID** | **Function** | ***St*-sRNA** | ***Pi*-sRNA** |
|---|---|---|---|
| PGSC0003DMG400016601 | Tospovirus resistance protein A | CACGAAUCCUUCCACGAGCUG | UGAAUCCUUCGGCUAUCCAU |
| PGSC0003DMG402016602 | Tospovirus resistance protein C | CACGAAUCCUUCCACGAGCUG | UGAAUCCUUCGGCUAUCCAU |
| PGSC0003DMG401022374 | Disease resistance protein (TIR class) | UCUCGAUCACGAAUCCUUCCA | UUGGAUUGAAGGGAGCUCUA |
| PGSC0003DMG400006573 | Tospovirus resistance protein A | CAAGGCACUCCAUACUUUCAG | UGAAUCCUUCGGCUAUCCAU |
| PGSC0003DMG400021986 | Late blight resistance protein Rpi-blb2 | UCCACUUCAGUCUUCAAAGU | UAAAGCAAGCAUGGCUUUAGGU |
| PGSC0003DMG400025512 | Late blight resistance protein Rpi-blb2 | UCCACUUCAGUCUUCAAAGU | UAAAGCAAGCAUGGCUUUAGGU |
| PGSC0003DMG400010895 | Late blight resistance protein Rpi-blb2 | UCCACUUCAGUCUUCAAAGU | UAAAGCAAGCAUGGCUUUAGGU |

**b,** Seven predicted *R* genes shared by the *St*-sRNA and *St*-miRNA pools

| **Target gene ID** | **Function** | ***St*-miRNA** | ***St*-sRNA** |
|---|---|---|---|
| PGSC0003DMG400020935 | Tir-nbs-lrr resistance protein | UUUUAGCAAGAGUUGUUUUCCC | UGUUAAGGAUUCUAAUUGGCU |
| PGSC0003DMG400006296 | Tospovirus resistance protein A | UUCCACAGCUUUCUUGAACUG | UUCCACAGCUUUCUUGAAC |
| PGSC0003DMG400025273 | Multidrug resistance pump | UGAAGCUGCCAGCAUGAUCUA | UGAAGCUGCCAGCAUGAUC |
| PGSC0003DMG400002357 | Bacterial spot disease resistance protein 4 | UUACCGAUUCCCCCCAUUCCAA | UCUGCUGCCAUUACCAAACCA |
| PGSC0003DMG400019627 | Disease resistance protein Gpa2 | UUUUAGCAAGAGUUGUUUUCCC | UAAAGCAAGCAUGGCUUUAGGU |
| PGSC0003DMG400004874 | Resistance protein PSH-RGH7 | UUUUAGCAAGAGUUGUUUUCCC | UUUCCUAUUCCACCCAUGC |
| PGSC0003DMG402026432 | NBS-coding resistance gene protein | UUUAGCAAGAGUUGUUUUCCC | UGUUAAGGAUUCUAAUUGGCU |

**c,** Two predicted *R* genes shared by the *St*-miRNA and *Pi*-sRNA pools.

| **Target gene ID** | **Function** | ***St*-miRNA** | ***Pi*-sRNA** |
|---|---|---|---|
| PGSC0003DMG400001081 | NBS-LRR protein | UCUUGCCAAUACCGCCCAUUCC | GGAGACCACAUCAAGAAGCUC |
| PGSC0003DMG400043400 | Disease resistance protein RGA2 | UUACCGAUUCCCCCCAUUCCAA | AUGUGUGACCAAACAUCAU |

**Table S4. Predicted miR8788 target transcripts in the *P. infestans* genome.**
Targets with expectation values up to 5.0 are shown.

**a,** miR8788-3p targets

| Target gene ID | Function |
|---|---|
| PITG_10391 | Unknown |
| PITG_12147 | Unknown |
| PITG_00350 | RPN7 |
| PITG_10532 | Unknown |
| PITG_21309 | Hist_deacetyl |
| PITG_01911 | Hist_deacetyl |
| PITG_13647 | XhoI |
| PITG_08884 | AhpC-TSA |
| PITG_08045 | RrnaAD |
| PITG_11501 | Pinin_SDK_memA |
| PITG_01437 | Med7 |
| PITG_08266[1] | ADH_zinc_N |
| PITG_03356 | EcKinase |
| PITG_07037 | LRR_6 |
| PITG_17709 | EXS |
| PITG_06886 | SpoU_methylase |

1. PITG_08266 is targeted at 3 sites (942-961, 1272-1291, 1602-1621)

**b,** miR8788-5p targets

| Target gene ID | Function |
|---|---|
| PITG_21918 | AAAP |
| PITG_20231 | AAAP |
| PITG_15406 | AAAP |
| PITG_17766 | AAAP |
| PITG_20230 | AAAP |
| PITG_20239 | ABC2_membrane_3 |
| PITG_19690 | Biotin_lipoyl |
